# Integrative Transcriptomic Analysis Reveals the Immune Mechanism for A CyHV-3-Resistant Common Carp Strain

**DOI:** 10.1101/2020.11.10.375915

**Authors:** Zhiying Jia, Nan Wu, Xiaona Jiang, Heng Li, Jiaxin Sun, Mijuan Shi, Chitao Li, Xuesong Hu, Yanlong Ge, Weidong Ye, Ying Tang, Junwei Shan, Yingyin Cheng, Xiao-Qin Xia, Lianyu Shi

## Abstract

Anti-disease breeding is becoming the most promising solution to cyprinid herpesvirus-3 (CyHV-3) infection, the major threat to common carp aquaculture. Virus challenging studies suggested that a breeding strain of common carp is resistant to this disease. This study illustrates the immune mechanisms involved in anti-CyHV-3 ability. An integrative analysis of the protein-coding genes and long non-coding RNAs (lncRNAs) using transcriptomic data was performed. Tissues from the head kidney of common carp were extracted at day 0 (the healthy control) and day 7 after CyHV-3 infection (the survivors), and used to analyze the transcriptome through both Illumina and PacBio sequencing. Following analysis of the GO terms and KEGG pathways involved, the immune-related terms and pathways were merged. In order to dig out details in the immune aspect, The DEGs were filtered using the current common carp immune gene library. Immune gene categories and their corresponding genes in different comparison groups were revealed. Also, the immunological Gene Ontology terms for lncRNA modulation were retained. The weighted gene co-XSexpression network analysis was used to reveal the regulation of immune genes by lncRNA. The results demonstrated that the breeding carp strain develops marked resistance to CyHV-3 through a specific innate immune mechanism. The featured biological processes were autophagy, phagocytosis, cytotoxicity, and virus blockage by lectins and MUC3. Moreover, the immune suppressive signals, such as suppression of IL21R on STAT3, PI3K mediated inhibition on inflammation by dopamine upon infection, as well as the inhibition of NLRC3 on STING during a steady state. Possible susceptible factors for CyHV-3, such as ITGB1, TLR18, and CCL4, were also revealed from the non-breeding strain. The results of this study also suggested that Nramp and PAI regulated by LncRNA could facilitate virus infection and proliferation for infected cells respectively, while T cell leukemia homeobox 3 (TLX3) as well as galectin 3 function by lncRNA may play a role in the resistance mechanism. Therefore, immune factors that are immunogenetically insensitive or susceptible to CyHV-3 infection have been revealed.

## INTRODUCTION

Cyprinid herpesvirus-3 (CyHV-3) infection is a major threat to common carp aquaculture (1), leading to widespread mortality and substantial economic loss. CyHV-3 is thought to cause death by weakening the host’s immune system, resulting in susceptibility to pathogenic microbes (1). In common carp, clinical signs of the disease develop rapidly and may induce morbidity and mortality within a period of 6 to 10 days following infection (2).

Carp that survive a primary infection with CyHV-3 can be resistant to future infection with this virus. Since latency and persistent carrying of CyHV-3 exist in carp (2–4), genetic backgrounds are crucial in developing an understanding resistance against the virus. Experimental infections of carp from pure lines or crosses have indicated the existence of genetic background of resistance by divergent survival rates (2). For example, a markedly higher expression of immune-related genes involved in pathogen recognition, complement activation, major histocompatibility complex class I (MHC I)-restricted antigen presentation, and the development of adaptive mucosal immunity was noted in the more resistant R3 line. Higher activation of CD8^+^ T cells was also observed (5). The diallelic cross of four European carp lines, including Polish ‘K’ and ‘R6’, Hungarian ‘R7’ and French ‘F’ also has been down in order to select the resistant fish, and then found that MH class II B genes of carp can have an effect on immunity against CyHV-3 infection (6).Additionally, carp strains of Asian origin, particularly Amur wild carp, were shown to be more resistant to CyHV-3 than strains originating from Europe, such as the Prerov scale carp or koi carp from a breed in the Czech Republic (7).

The immune response of carp to CyHV-3 involves both innate and adaptive aspects with the outcome of the disease largely depending on whether the balance is tipped in favor of the host’s immune response or virus’s evasion strategy (2). In general, transcriptomic analysis has revealed that three immune-related pathways, namely the mitogen-activated protein kinase (MAPK) signaling pathway, the innate immune response, and the cytokine-mediated signaling pathway, were highly involved in the infection with CyHV-3 (8). In red common carp × koi, the expression of interleukin 12 (IL12) p35, interferon (IFN) αβ, and toll-like receptor 9 (TLR9) may provide potential genes related to resistance against KHV (another term for CyHV-3) (9). However, the magnitude of type I IFN response did not correlate with a higher resistance in CyHV-3-infected carp, during the challenge test among different strains, although CyHV-3 infection can induce type I IFNs (7, 10). Regarding the innate resistance in carp, the mapped CyHV-3 survival quantitative trait loci have been reported mainly in *IL10* and *MHC II* (11). Recently, by quantitative trait locus mapping and genome-wide association study, *tumor necrosis factor-alpha* (*TNFA*), *hypoxia inducible factor 1 subunit alpha* (*HIF1A*), *galectin-8* (*LGALS8*), *rootletin*, and *paladin*, have also been related to resistance against CyHV-3 (12). Adaptive immunity through both cytotoxicity and immunoglobulin (Ig) secretion may be involved in resistance. Matthew et al. revealed that, in the anterior kidney, Ig secretion plays an important role in the resistance during the persistent infection or reactivation phases of CyHV-3 (1). In addition, CyHV-3 profoundly influences the expression of host miRNA, although the regulation of immune processes by miRNA in the clinical and latent phases differs (4). This suggests an important role of non-coding RNAs in anti-CyHV-3 immunity.

The first transcriptional analysis of carp anterior kidney mainly pointed out the important role of humoral immune responses, especially those related to immunoglobulin (1). Spleen transcriptomic analyses comparing the susceptible and resistant common carp revealed that the susceptible fish elicited a typical anti-viral interferon response, while the upregulated IL-8 attracted innate immune cells and related response may play an essential role in resistant fish (13).

German mirror carp selection G_4_ is a strain of common carp cultivated widely in China, yet with a high mortality caused by CyHV-3 virus. Based on our previous study, a strain of common carp from German mirror carp selection G_4_ has already shown a higher survival rate after breeding for three generations. Among fish from the G_3_ generation, 1000 individuals were genotyped by four SNP loci, including carp065309, carp070076, carp183811 and carp160380, and then four main groups, with genotypes of GG/GT/GG/AA, GG/TT/TT/AA, AG/TT/AG/AT and GG/GG/GG/AA, whose survival rates were 89.9%, 94.7%, 88.0% and 92.7% respectively, were propagated (14). While the unselected mirror carp were 62.6%. Since decreased viral load in tissues directly indicate the resistance to CyHV-3 (15), the fact that it has displayed a better immunological index as well as a reduced virus load in immune organs, such as the kidneys and spleen (16, 17), strongly suggests the resistance to CyHV-3 in current used breeding strain. In detail, acid phosphatase in the spleen, glutathione and total antioxidation capacity in the kidney, lysozyme and immunoglobulin M in the serum, and alkaline phosphatase in the spleen and kidney also showed significant differences between G_1_ and G_3_ or G_1_ and G_2_/G_3_. Meanwhile, the survival rate after CyHV-3 challenge increased generation by generation. The strong resistance to CyHV-3 has been stable for G_3_ (16). Recently, Sun has revealed that virus genes *TK* and *ORF72* in the G_4_ were expressed at levels significantly lower than those in the non-breeding strain (17). Thus, the resistance to CyHV-3 in G_4_ was strongly deduced.

However, there is a lack of systemic studies focusing on a detailed network of the immune system for the anti-CyHV-3 immune mechanism in this resistant strain. The third-generation sequencing method, such as PacBio, can decode the genetic sequences that are markedly longer compared with the second-generation method, such as Illumina (18). Therefore, a higher quality assembly of transcripts, including both mRNA and non-coding RNAs, may provide a more detailed understanding of the mechanism of genetic resistance.

In this study, in order to reveal the immune mechanisms involved in anti-CyHV3 breeding for common carp, integrative transcriptomic analysis was performed for both mRNA and long non-coding RNA (lncRNA) during the CyHV-3 challenge, using strategy as shown in Fig. 1. Immune-related transcripts, from both survivors and healthy controls of either breeding or non-breeding strains, were analyzed. By comparing data from different groups for different strains obtained through both Illumina (New England Biolabs, Ipswich, MA, USA) and Pacific Biosciences (PacBio) sequencing, both immune-related differentially expressed genes (DEG) and lncRNA have been revealed for the Kyoto Encyclopedia of Genes and Genomes (KEGG) pathways and/or Gene Ontology (GO) terms involved. The immune gene-related lncRNA was also explored. This study may shed some light on fish breeding for virus-resistant strains, and facilitate disease control after CyHV-3 infection.

**Figure 1.**
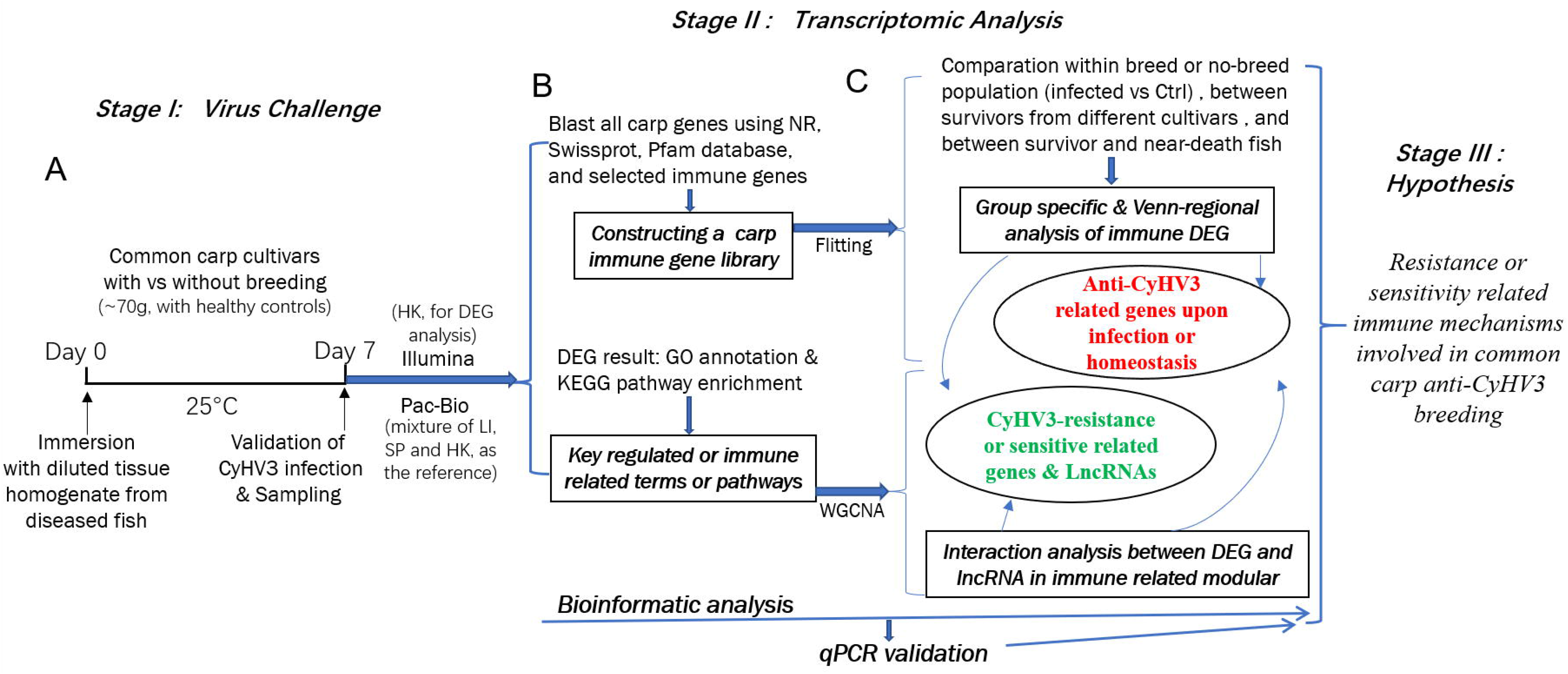
The strategy of integrative transcriptomic analysis for revealing immune mechanisms involved for resistance or sensitivity for current anti-CyHV3 breeding in common carp. During stage I, the disease modelling for CyHV3 infection was established using immersion of virus containing tissue homogenate, and then validated by checking the expression of virus genes. The transcriptomic sampling was done at day 7 post infection for sequencing using both Illumina and Pac-Bio platforms. In stage II, the bioinformatic analysis was carried out for the transcriptomic data. On one hand, DEG was generated from five comparison groups, including intra-strain comparison groups A (survivors vs. controls in the breeding strain) and B (survivors vs. controls in the non-breeding strain), inter-strain comparison groups C (the comparison between survivors at day 7 for the two strains) and D (the comparison between control fish at day 0 for the two strains), and group E. The vital genes for survival were investigated through group E, in which the genes, without significant differences between the survivors from the breeding and non-breeding strains, were compared with the genes from severely sick fish, which swam very slowly, or even floated on the water. The revealed DEGs were annotated by GO terms and KEGG pathways. On the other hand, a common carp immune gene library was constructed for further immune related analysis. Additionally, in order to reveal the LncRNA regulation, WGCNA was used to reveal the interaction between differential expressed transcripts and LncRNA in immune related module. Thereafter, the resistance or sensitivity related immune mechanisms involved in common carp anti-CyHV3 breeding could be enlightened accordingly.

## MATERIALS AND METHODS

### Ethics Statement

All procedures involving animals in this study were conducted according to the guidelines for the care and use of laboratory animals of Heilongjiang River Fisheries Research Institute, Chinese Academy of Fishery Sciences (Harbin, China). The studies involving animals were reviewed and approved by the Committee for the Welfare and Ethics of Laboratory Animals of Heilongjiang River Fisheries Research Institute, Chinese Academy of Fishery Sciences.

### Animals and Virus Challenge

German mirror carp selection G_4_ used in this study were obtained from Kuandian Research Base of Heilongjiang River Fisheries Research Institute (Liaoning, China). The breeding strain was the fourth generation after the selection of resistance to CyHV-3 (17). Two G4 groups with survival rates of 94.7% and 92.7% were mixed with the ratio 1:1 and used as the experimental populations in this study. Both the breeding strain G_4_ and non-breeding strain were used for the virus challenge. Since CyHV-3 has a mucosal route of infection mainly via the skin rather than the gut (2), this study induced infection with CyHV-3 by adding the homogenate solution of internal organs from sick fish into the water of the tanks following the previously published methods (16, 17, 19). The homogenate solution was prepared using organs from 10 severely sick fish, which swam very slowly, or even floated on the water. Before using the homogenate to infect fish, the head kidney tissues from those sick fish were checked for the CyHV-3 infection by PCR of virus genes *TK* and *Sph* following the method described in the industry standard SC/T 7212.1-2011 (20). To separate the control and experimental groups from either the breeding or non-breeding strains, four tanks (1.6 m × 1.2 m × 0.6 m) containing healthy juvenile common carp (~70 g; N=300 fish per tank), were used. The water temperature during the experiment was maintained at 25±1°C.

### Sampling and pathological analysis

A selection of common carp, from either the breeding or non-breeding strains, were sacrificed. Head kidney samples were obtained at days 0 and 7 after challenge, representing the control and survivor carp. The samples (N=3) were collected immediately and soaked in 10 volumes of RNAlater (Qiagen, Hilden, Germany), for sequencing using Illumina (New England Biolabs). To better sequence and generate the lncRNAs, PacBio sequencing was applied to analyze the mixture of liver, spleen, and kidney, with two replicates. The number of dead fish during the experiment was counted daily to calculate the mortality rate. Meanwhile, in order to analyze the swelling degree of inner organs, the proportion of trunk kidney to the whole trunk area was calculated by ImageJ (https://imagej.en.softonic.com/). In detail, the area of both trunk kidney and trunk regions were selected and measured, and the percentages between them was calculated based on 8 survivors of breeding or non-breeding strain. T-test was used to judge the significance of the difference.

### RNA Extraction

RNA was isolated using the AllPrep DNA/RNA FFPE Kit (Qiagen, Hilden, Germany), according to the instructions provided by the manufacturer. RNA degradation and contamination were monitored on 1% agarose gels. RNA purity was checked using the NanoPhotometer spectrophotometer (Implen Inc., Westlake Village, CA, USA). RNA concentration was measured using the Qubit RNA Assay Kit in a Qubit 2.0 Flurometer (Life Technologies, Carlsbad, CA, USA). RNA integrity was assessed using the RNA Nano 6000 Assay Kit of the Agilent Bioanalyzer 2100 system (Agilent Technologies, Santa Clara, CA, USA).

### Library Preparation and Sequencing for Transcriptomic Analysis

Optimized RNA-Seq strategies, including both PacBio and Illumina (New England Biolabs) sequencing (21), were used to more precisely resolve the sequence of transcripts. Firstly, the total RNA isolated from the head kidney was used to construct the cDNA library. Subsequently, the library was sequenced on a PacBio RS II platform (Biomarker Technologies, Beijing, China). For the PacBio Long Read Processing, raw reads were processed into error-corrected reads of insert using Iso-seq pipeline with minFullPass = 0 and minPredictedAccuracy = 0.90. Next, full-length, non-chemiric transcripts were determined by searching for the polyA tail signal and the 5’ and 3’ cDNA primers in reads of insert. Iterative Clustering for Error Correction was used to obtain consensus isoforms, and full-length consensus sequences from Iterative Clustering for Error Correction were polished using Quiver. High-quality transcripts with post-correction accuracy of >99% were retained for further analysis.

The illustration of transcripts obtained from the results of PacBio could provide reference transcriptional sequences for the assembly of Illumine sequencing data, to improve sequencing quality. Therefore, Illumina (New England Biolabs) sequencing was performed on all head kidney samples. The procedure used for the preparation of the gene library and sequencing of the transcriptome followed previously published methods (22). Briefly, sequencing libraries were generated using the NEBNext UltraTM RNA Library Prep Kit for Illumina (New England Biolabs), and the library quality was assessed on the Agilent Bioanalyzer 2100 system. The library preparations were sequenced on an Illumina platform, and 150 bp paired-end reads were generated.

### Annotation and Functional Analysis of Transcripts by Public Databases

All reads in the transcriptome data were mapped on the common carp genome (https://asia.ensembl.org/Cyprinus_carpio_german_mirror/Info/Index) (23) for annotation. The data from the Illumina platform were also used to check and replace errors in the data from the PacBio platform. Both the annotation of genes and lncRNAs were subsequently generated. The protein coding transcripts were annotated by NR, swissprot and Pfam database. DEGs were detected from five comparison groups, including intra-strain comparison groups A (survivors vs. controls in the breeding strain) and B (survivors vs. controls in the non-breeding strain), inter-strain comparison groups C (the comparison between survivors at day 7 for the two strains) and D (the comparison between control fish at day 0 for the two strains), and group E. The vital genes for survival were investigated through group E, in which the genes without significant differences (*p* > 0.05) between the survivors from the breeding and non-breeding strains were compared with the transcripts from severely sick fish. The revealed DEGs were annotated by GO terms and KEGG pathways, following a previously published protocol (22, 24). In brief, functional annotation and the classification of genes were determined by both employing local genes blasts against GO Consortium (http://geneontology.org/) and KEGG (https://www.kegg.jp/kegg/pathway.html). Enrichment of the KEGG pathways was carried out for both upregulated and downregulated genes for all comparison groups. Then in order to demonstrate the immune DEGs involved pathways more clearly, the gene list of the current construct common carp immune gene library was used as follows. In addition, the LncRNAs were also annotated by GO terms.

### Construction of the Common Carp Immune Gene Library

The common carp immune library was constructed by following our previously published method, which was applied to grass carp (22) and tilapia (24), with some adjustment for viral infection-related immune genes. The modifications were based on gene information obtained by blasting each sequence to databases, including NCBI NR database, as well as Swiss-Prot and Pfam databases. The common carp immune gene library contained information for immune genes at two levels. For the first level, nine categories of immune processes, namely “acute phase reactions”, “pattern recognition”, “antigen processing and regulators”, “complement system”, “inflammatory cytokines and receptors”, “adapters, effectors and signal transducers”, “innate immune cells related”, “T/B cell antigen activation”, and “other genes related to immune response”, were proposed. Subsequently, many categories of immune genes for each immune process (detailed in Table S1) were applied for the second level. The library was used to filter transcriptome data and obtain details of the immune processes and particular immune genes for each comparison group, during the GO term and KEGG pathway enrichment.

### Statistics Analysis

The DEG were generated by comparing the RPKM (Reads Per Kilobase of transcript, per Million mapped reads) using the DESeq2 R package (1.16.1). The resulting *p* values were adjusted using the Benjamini and Hochberg approach for controlling the false discovery rate. Genes with an adjusted *p* < 0.05 found by DESeq2 were assigned as being differentially expressed. The following bioinformatics analysis was performed to select immune-related transcripts from the common carp immune gene library, and construct the barplots for the major immune processes and immune categories. The t-test was used to assess differences, with a false discovery rate adjusted *p* < 0.05. Qualitative comparisons were performed between samples by counting the number of DEG. The data were rearranged in Microsoft EXCEL, and applied to plot charts by ggplot2 (2.2.1) using R language.

### Correlation Analysis between LncRNA and Genes Involved in Immune-related GO Terms

The weighted gene co-expression network analysis (WGCNA) was performed using the R package “WGCNA” (25). Specifically, all genes with an expression variance ranked in the top 75 percentile of the data set were retained (26). The R package WGCNA was used to construct the weighted gene co-expression network (27). A matrix of signed Pearson correlations between all gene pairs was computed, and the transformed matrix was used as input for linkage hierarchical clustering. Genes with similar expression patterns were clustered together as one module. Subsequently, using R package clusterProfiler (28), enriched GO terms for lncRNA-related protein-coding genes were generated for the gene list of every module. The *p* values of enriched GO terms were produced from Kolmogorov-Smirnov test. In order to elucidate the detail lncRNA-mRNA network, the immune-related GO term containing module were selected, and the relationship of involved genes and related lncRNAs were shown by the cystoscope software. In addition, the immune genes in immune related module were classified into different comparison groups.

### Reverse Transcription Polymerase Chain Reaction (RT-PCR)

The mRNA samples used for transcriptome sequencing were also subjected to real-time quantitative RT-PCR validation (n = 3), following a previously published protocol [14, 16]. After RNA isolation, reverse transcription and quantitative PCR (qPCR) experiments were carried out for 11 genes. Gene expression levels were determined using TaqMan-based qPCR (Thermo Fisher Scientific, Waltham, MA, USA), which was performed on the RotorGene system (Qiagen, Hilden, Germany). The total RNA was reverse transcribed, and 60 ng were used for the One Taq RT PCR Kit (New England Biolabs) according to the instructions provided by the manufacturer for mRNA quantification and reverse transcription (Thermo Fisher Scientific, Waltham, MA, USA). For mRNA quantification, the resulting cDNA was pre-amplified using a TaqMan PreAmp Master Mix Kit (Thermo Fisher Scientific, Waltham, MA, USA). The relative levels of mRNA were determined from diluted cDNA and pre-amplified cDNA using qPCR and appropriate TaqMan probes. The expression levels were calculated after an efficiency correction (based on the results of a serial dilution) relative to those of endogenous controls.

## RESULTS

### Virus identification, Mortality Rate and Pathological Appearance

The successful infection of CyHV-3 has been validated using PCR, in order to check the expression of virus genes (*TK* and *Sph*), in randomly sampled severely sick fish (N=10) at day 7 after challenge (Figure 2A). Further, to reveal the mortality, the number of dead fish was recorded from day 1 to day 15 after challenge (Table S1), and then the mortality rate was calculated for all groups (Table S2). For the challenged non-breeding strain (Figure 1B), the mortality rate increased daily, except for a decrease observed on day 5. Mortality peaked at day 7 (69%) and decreased to a markedly low level thereafter. For the challenged breeding strain (Figure 2B), the mortality rate remained at a markable low level, with the highest value (11%) recorded at day 7. The mortality rates of the two challenged groups were very significantly different (*p* < 0.01). The number of dead fish for each day is listed in Table 1 for the breeding or non-breeding strain, while in the two unchallenged controls, there were no dead fish (data not shown). Compared with the organs in survivors of the breeding strain, pathological swelling and hyperemia of the internal organs (particularly the kidney) was obvious in survivors from the non-breeding strain on day 7 (Figure 2C). In detail, the proportion of kidney area to the whole trunk area in the breeding strain was significantly different from that of the non-breeding strain (*p* < 0.01) (Figure 2D). The reduced immune organ swelling was reflected by the ratio of 0.44 comparing the proportions between the breeding and non-breeding strains (Table S2).

**Figure 2.**
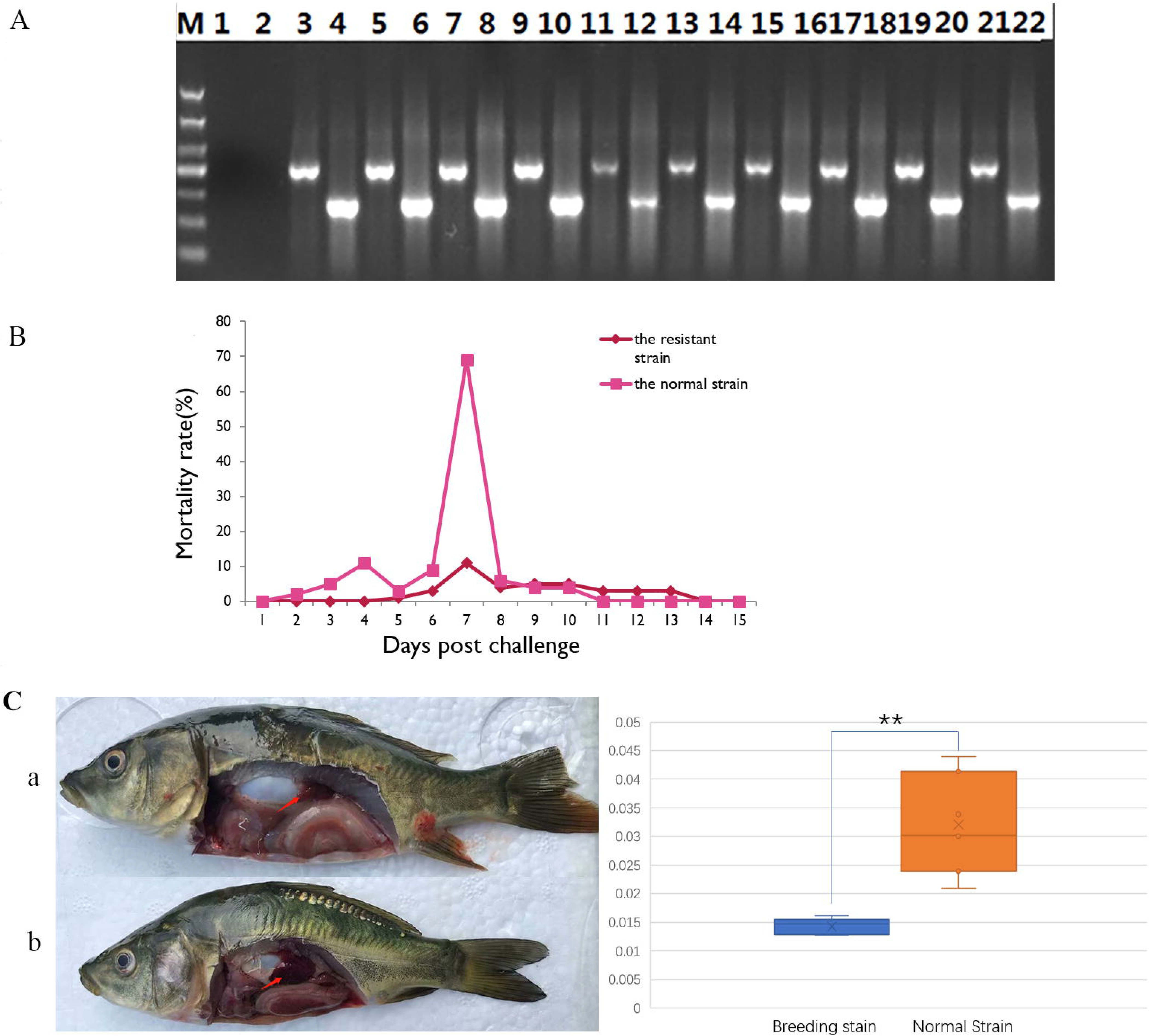
Differential appearance of general mortality and pathology between fish from the breeding and non-breeding strains. (A) The PCR validation of the CyHV-3 virus genes *TK* and *Sph*. The lane 1-2 represents the negative control, and afterwards lane 3-22 represents the result for tested 10 virus infected fish. The lanes of odd and even numbers showed the PCR result of *TK* and *Sph* genes, respectively. M is the abbreviation of “marker”. **(B)** Comparison of daily mortality between the breeding and non-breeding strains. The mortality was monitored within 15 days post challenge. N=300 per group. **(C)** The degree of swelling trunk kidney was limited in the survivors from the breeding strain (a) compared with the markedly enlarged trunk kidney observed in fish from the non-breeding strain (b). The arrow indicates the trunk kidney region.

**Table 1.** The number of dead fish for each day during the CyHV-3 challenging for breeding or non-breeding stain

| Stain | D1 | D2 | D3 | D4 | D5 | D6 | D7 | D8 | D9 | D10 | D11 | D12 | D13 | D14 | D15 |
| --- | --- | --- | --- | --- | --- | --- | --- | --- | --- | --- | --- | --- | --- | --- | --- |
| Breeding | 0 | 0 | 0 | 0 | 1 | 3 | 11 | 4 | 5 | 5 | 3 | 3 | 3 | 0 | 0 |
| Non-breeding | 0 | 2 | 5 | 11 | 3 | 9 | 69 | 6 | 4 | 4 | 0 | 0 | 0 | 0 | 0 |

### Quality and Validation of Transcriptomic Data

Sequencing was performed with both the PacBio RS II and Illumina platforms to analyze the gene information of the common carp. Through PacBio sequencing, 19.49 G data were obtained. Of the 258,346 ccs, 79.79% were full-length sequences, and 17,769 polished high-quality isoforms were also revealed (Table S3, sheet 1). Meanwhile, the Illumina data were also characterized by high quality, and the information for each sample is provided in Table S3 (sheet 2). In order to validate the transcriptomic data, qPCR was conducted. A total of 24 reactions were performed to validate the transcripts of 11 genes, most of which were immune genes. The correlation analysis showed that fold changes between transcriptome and qPCR results correlated well (R^2^=0.929718564). The gene ID, annotation, primers, and fold-change information are listed in Table S4.

### GO Analysis of DEGs among Tested Fish

According to the enriched general GO terms (Figure 3) of immune related DEGs, there are “immune system process” and “antioxidant activity” for all comparison groups, “chemoattractant activity” only in comparison group A, “rhythmic process” only in comparison group B, “cell killing”in both comparison groups A and C, and “virion” in both comparison groups B and C. The detail significant enriched GO terms involved in biological processes (BP), cellular components (CC), and molecular functions (MF) were detailed in Table S5-S9 for all comparison groups.

**Figure 3.**
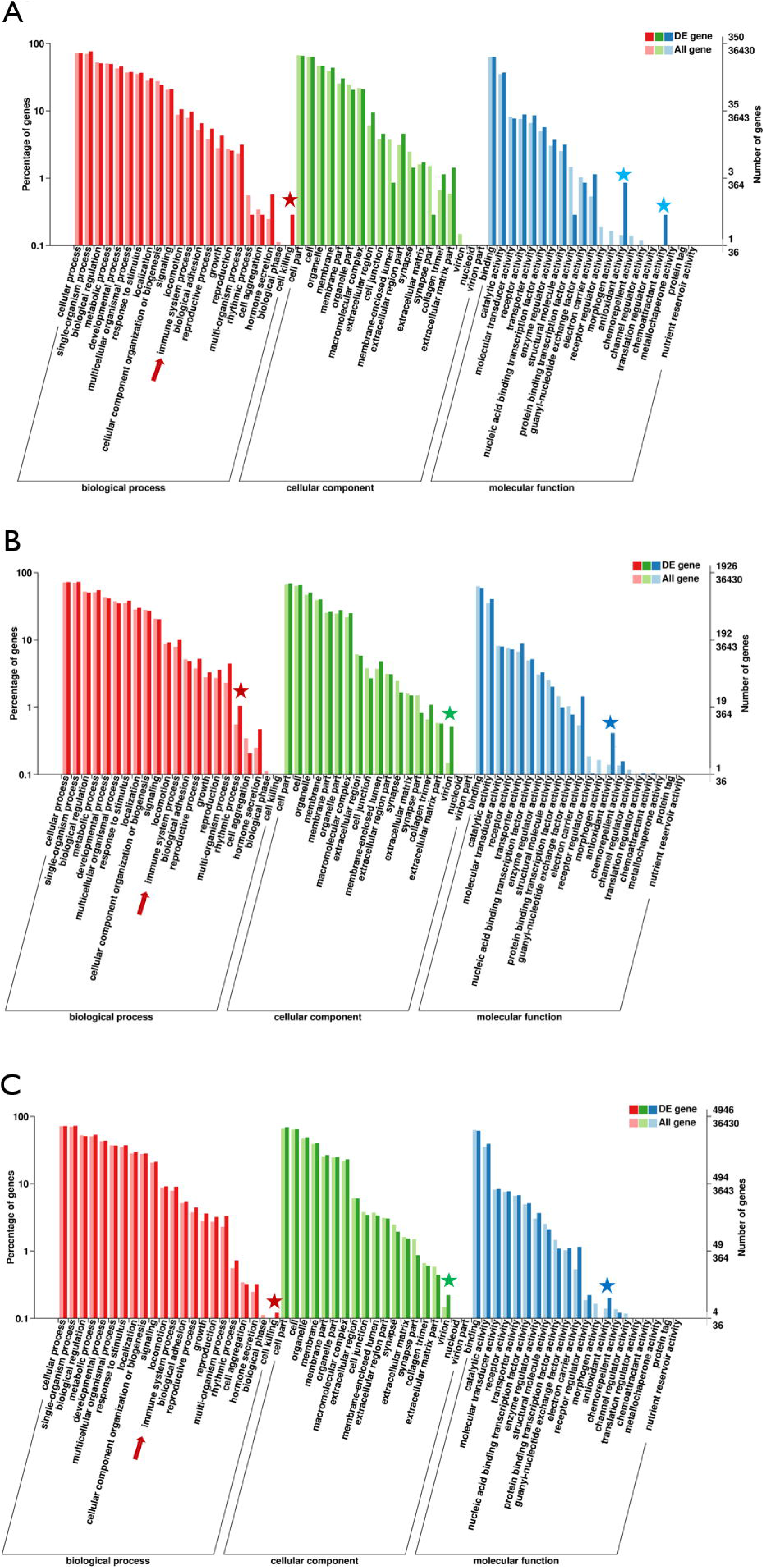
GO (Gene Ontology) terms of comparison groups A, B, and C. The different colors of bar indicate the comparison of the GO terms between all genes and immune related genes. In detail, red and pink bars represent all genes and immune related genes respectively for the biological process (BP), green and light green bars represent those for the cellular component (CC), blue and light blue bars represent those for the molecular function (MF). The immune-related terms are labeled with stars.

The featured BP terms (*p* < 0.01) include “phagocytosis”, “positive regulation of immune system process”, “positive regulation of I-kappa B kunase/NF-kappaB signaling”, “integrin-mediated signaling pathway”, “regulation of B cell activation”, “macrophage activation”, “lymphocyte mediated immunity” in immune aspect in group A. In group B, the BP terms include “response to lipopolysaccharide”, “regulation of granulocyte differentiation”, “toll-like receptor signaling pathway”, “positive regulation of immune system process”, “macrophage differentiation”, “phagocytosis”, “engulfment”, “lymphocyte activation”, “integrin-mediated signaling pathway” and “leukocyte activation”. In group C, the BP terms include “Response to lipopolysaccharide”, “negative regulation of B cell apoptotic process”, “positive regulation of TOR signaling”, “macrophage differentiation”, “endosome to lysosome transport”, “regulation of granulocyte differentiation”, “toll-like receptor signaling pathway”, “defense response to fungus”, “phagocytosis”, “engulfment”, “T cell proliferation”, “mast cell activation”, “myeloid cell activation involved in immune response”, “regulation of T cell differentiation in thymus”, “phagosome maturation”, and “positive regulation of T cell activation”. In group D, the BP terms include “Phagocytosis”, “positive regulation of immune system process”, “positive regulation of I-kappaB kinase/NF-kappaB signaling”, “integrin mediated signaling pathway”, “regulation of B cell activation”, “lymphocyte mediated immunity”, “macrophage activation”, and “response to lipopolysaccharide”. In group D, the BP terms include “Phagocytosis”, “positive regulation of cell migration”, “positive regulation of I-kappaB kinase/NF-kappaB signaling”, “positive regulation of myeloid leukocyte differentiation”, “integrin-mediated signaling pathway”, “regulation of B cell activation”, “leukocyte activation”, “regulation of immune response”, and “macrophage activation”.

Meanwhile, several immune related BP terms (*p* < 0.01) were also revealed, including “positive regulation of cell migration” in group A; “activation of MAPK activity”, “regulation of protein ubiquitination involved in ubiquitin-dependent protein catabolic process”, “oxication-reduction process”, “positive regulation of cell migration” in group B; “activation of MAPK activity”, “dopaminergic neuron differentiation” in group C. Besides, the CC terms (*p* < 0.01) includes “Pre-autophagosomal structure membrane” in group A, B, D and E, and the MF terms (*p* < 0.01) includes “NF-KappaB binding” and “lysozyme activity” in group B; “cytokine activity” in group C; “Lysozyme activity”, “MAP kinase activity” and “ubiquitin-like protein binding” in group D.

### KEGG Analysis of DEGs among Tested Fish

Compared with group A (survivors vs control in the breeding strain), there were markedly more genes involved in immune and related pathways in groups B (survivors vs control in the non-breeding stain). Current revealed immune pathways in group A were “phagosome”, “regulation of autophagy”, “ubiquitin mediated proteolysis”, and “plant-pathogen interaction” (Figure 4Aa). Meanwhile, there are four immune pathways, including “endocytosis”, “phagosome”, “FoxO signaling pathway”, “ubiquitin mediated proteolysis”, “DNA replication”, “fructose and mannose metabolism”, “oxidative phosphorylation”, and “plant-pathogen interaction” in group B (Figure 4Ab). To get better understanding of the different survival mechanism, the pathways revealed in group C (DEG, obtained from comparing survivors from the non-breeding to breeding strain) includes “endocytosis”, “phagosome”, “ubiquitin mediated proteolysis”, “DNA replication”, “fructose and mannose metabolism”, and “oxidative phosphorylation” in group C (Figure 4Ac, left).

**Figure 4.**
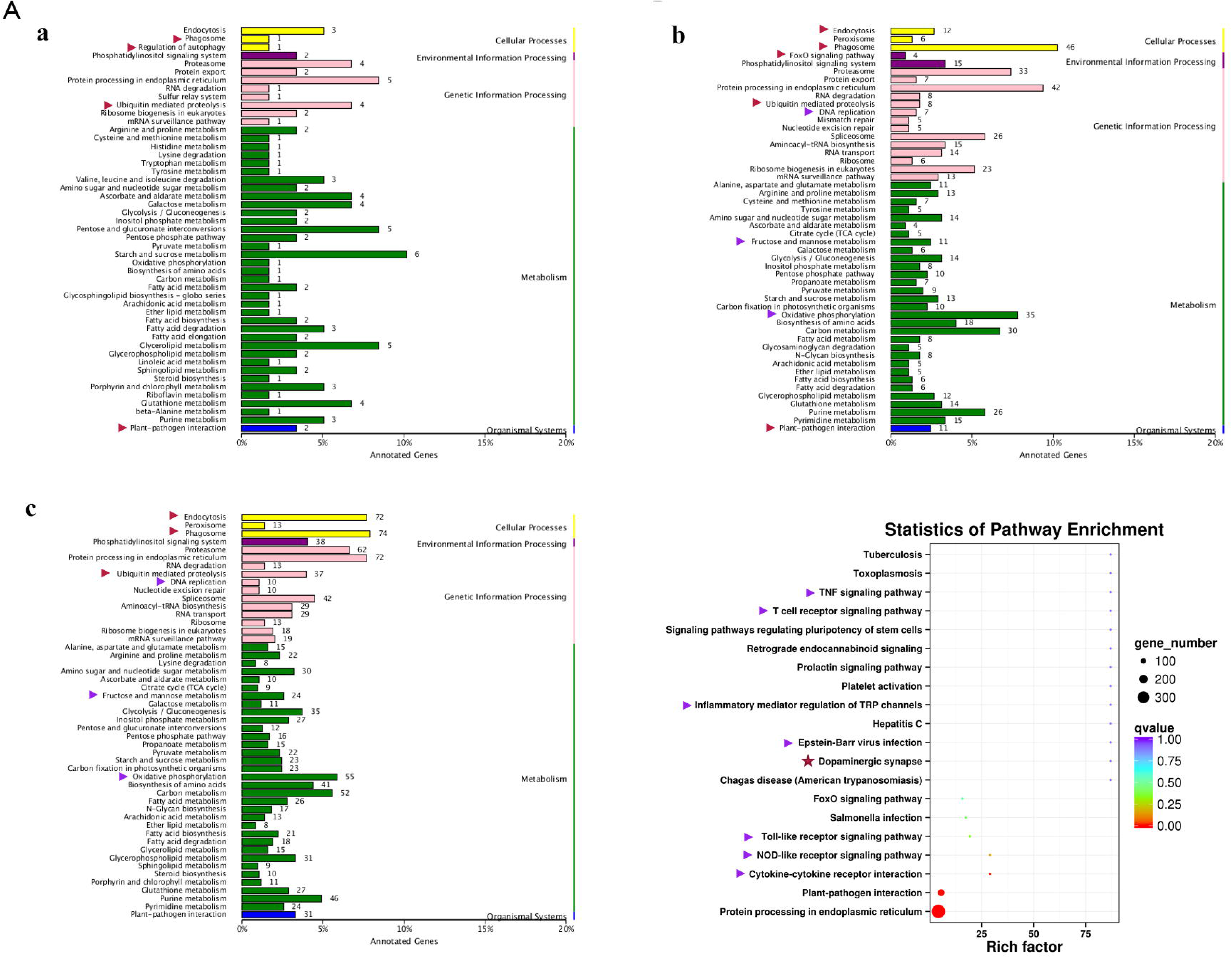
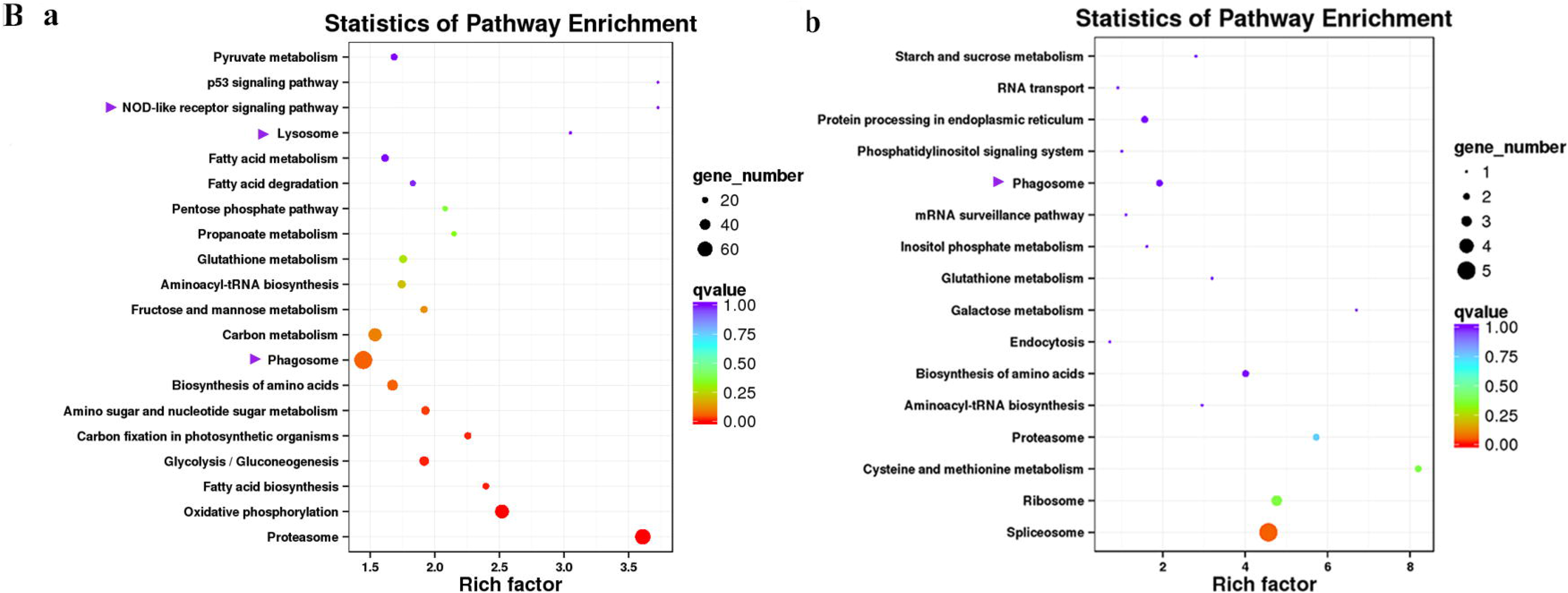
KEGG (Kyoto Encyclopedia of Genes and Genomes) pathways revealed for comparison groups A-E. In part A, the enriched pathways for comparison group A (a), B (b) and C(c) were showed mainly in bar plots. In addition, for the comparison group C, the babble plot of pathways for immune related DEG was also given in the right in order to illustrate the rich factor. In part B, the pathways of immune related DEG for both comparison group D(a), which compared controls of both stains, and group E(b), which has compared similar genes of survivors from both strains with protein coding transcripts of severely sick fish, were demonstrated in babble plots. The immune-related terms are labeled with arrows. Red and purple arrows indicate the immune and related pathways representatively.

In order to better illustrate the immune and related pathways more clearly, the filtering of DEG was conducted using the gene list of the common carp immune gene library (Table S1), and then bubble chart was used to clarify both the enrichment factors and gene numbers for revealed immune pathways. For group C, which reflected differential surviving mechanism (Figure 4A (c, right)), the revealed immune pathways include “TNF signaling pathway”, “T cell receptor signaling pathway”, “inflammatory mediator regulation of TRP channels”, “Epstein-Barr virus infection”, “Toll-like receptor signaling pathway”, “NOD-like receptor signaling pathway” and “cytokine-cytokine receptor interaction”. In addition, “dopaminergic synapse” was also found. While, groups D (DEG, obtained from compared controls of both stains) and E (DEG obtained from comparing similar genes of survivors from both strains with protein coding transcripts of severely sick fish), revealed the difference and similarity for maintaining basic homeostasis, respectively. The most enriched immune pathways were “p53 signaling pathway”, “NOD-like receptor signaling pathway”, “lysosome”, “oxidative phosphorylation”, and “proteasome” in group D (Figure 4Ba), yet only “phagosome” in group E (Figure 4Bb).

### Classification of DEG from Different Comparison Groups into Immune Process and Immune Gene Category

The current construction of the common carp immune gene library (Table S4) was used to classify the revealed immune transcripts and refine the involved immune processes and immune gene categories. The details for comparison groups A–E were provided in Tables S5–S9, respectively. At the level of immune process, in group A (Figure 5A, Table 2A and Table S5), most immune mRNAs were upregulated in the immune processes of “pattern recognition”, “inflammatory cytokines and receptors”, and “T/B cell antigen activation”, while downregulated in the immune process of “inflammatory cytokines and receptors”, “complement system”, “adapters, effectors and signal transducers”, and “pattern recognition”. Interestingly, there were no genes in “antigen processing and regulators” for group A. In addition, there were only downregulated genes in “acute phase reactions”, as well as only upregulated genes in “innate immune cells related” in group A. In group B (Figure 5A, Table 2A and Table S6), immune genes were upregulated in the immune processes, such as “other genes related to immune response”, “inflammatory cytokines and receptors”, and “adapters, effectors and signal transducers”. Meanwhile, the downregulated genes were involved in the immune processes, such as “pattern recognition”, “inflammatory cytokines and receptors” and “other genes related immune response”. The largest number of immune mRNAs was observed in group C (Figure 5A, Table 2A, and Table S7). The upregulated genes were mainly involved in “inflammatory cytokines and receptors”, “other genes related to immune response”, “T/B cell antigen activation”, “pattern recognition”, and “adapters, effectors and signal transducers”, whereas the downregulated genes were mainly involved in “other genes related to immune response”, “adapters, effectors and signal transducers”, “inflammatory cytokines and receptors”, “T/B cell antigen activation”, “pattern recognition”, and “antigen processing and regulators”. Meanwhile, in group D (Figure 5A, Table 2B and Table S8), the only upregulated genes were reported in “antigen processing and regulators” and “T/B cell antigen activation”, while downregulated genes were noted in “complement system”. The upregulated genes in group D were mostly involved in “adapters, effectors and signal transducers”, and few in “inflammatory cytokines and receptors”, whereas the downregulated genes were mostly involved in “inflammatory cytokines and receptors”. In group E (Figure 5B, Table 2B and Table S9), among revealed six processes, “T/B cell antigen activation” and “other genes related to immune response” exhibited the greatest DEG number. To illustrate the immune process, the involved immune gene categories of the immune-related DEG in all comparisons were listed in Table 2.

**Figure 5.**
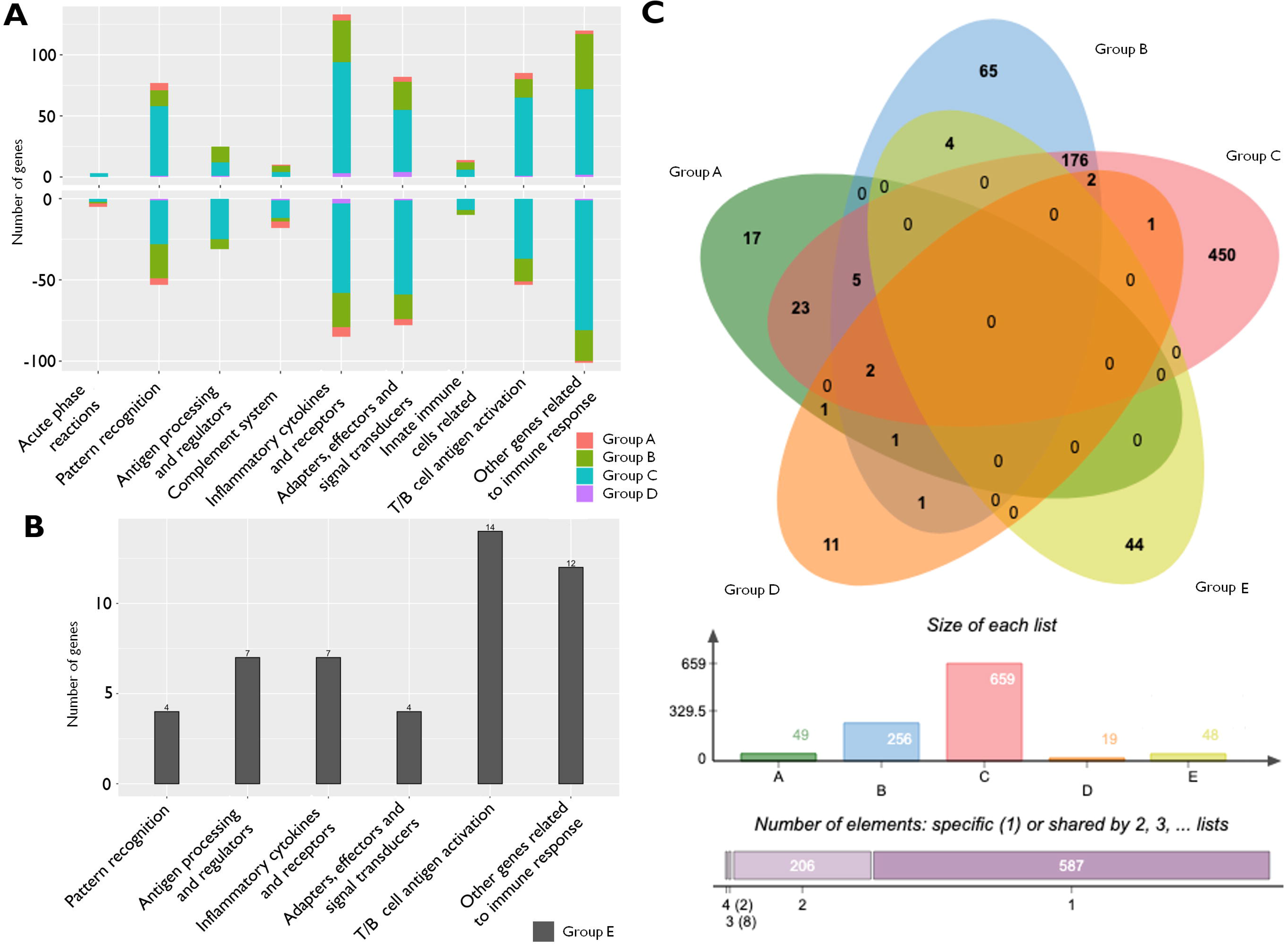
The analysis of immune processes of DEG among different comparison groups. (A) The barplots of gene number for the involved immune processes of up or down regulated DEGs in groups A-D; (B) The barplot of gene number for the involved immune processes in group E; (C) The Venn diagram of gene number for groups A-E. DEG, differentially expressed gene.

**Table 2A.**
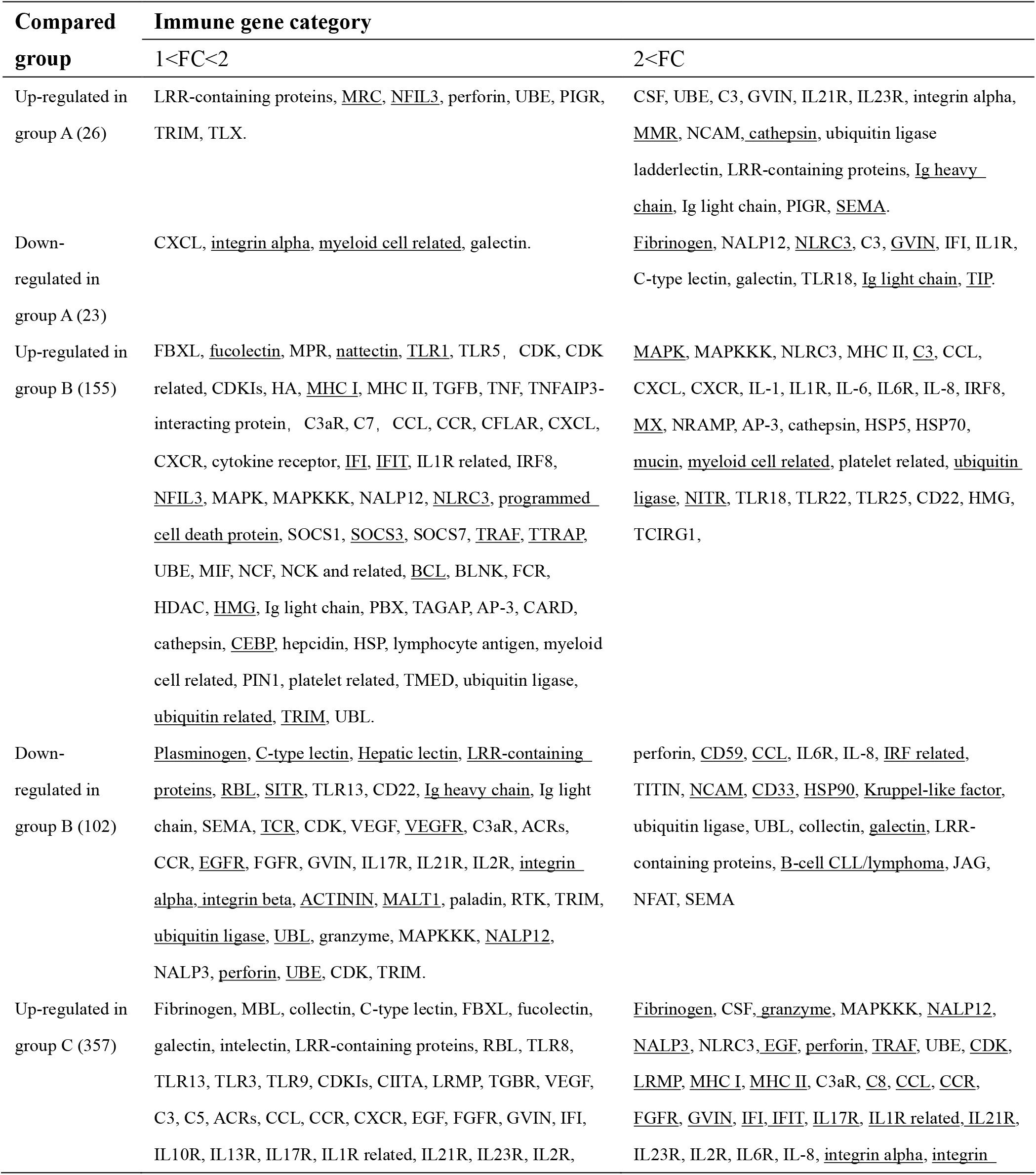
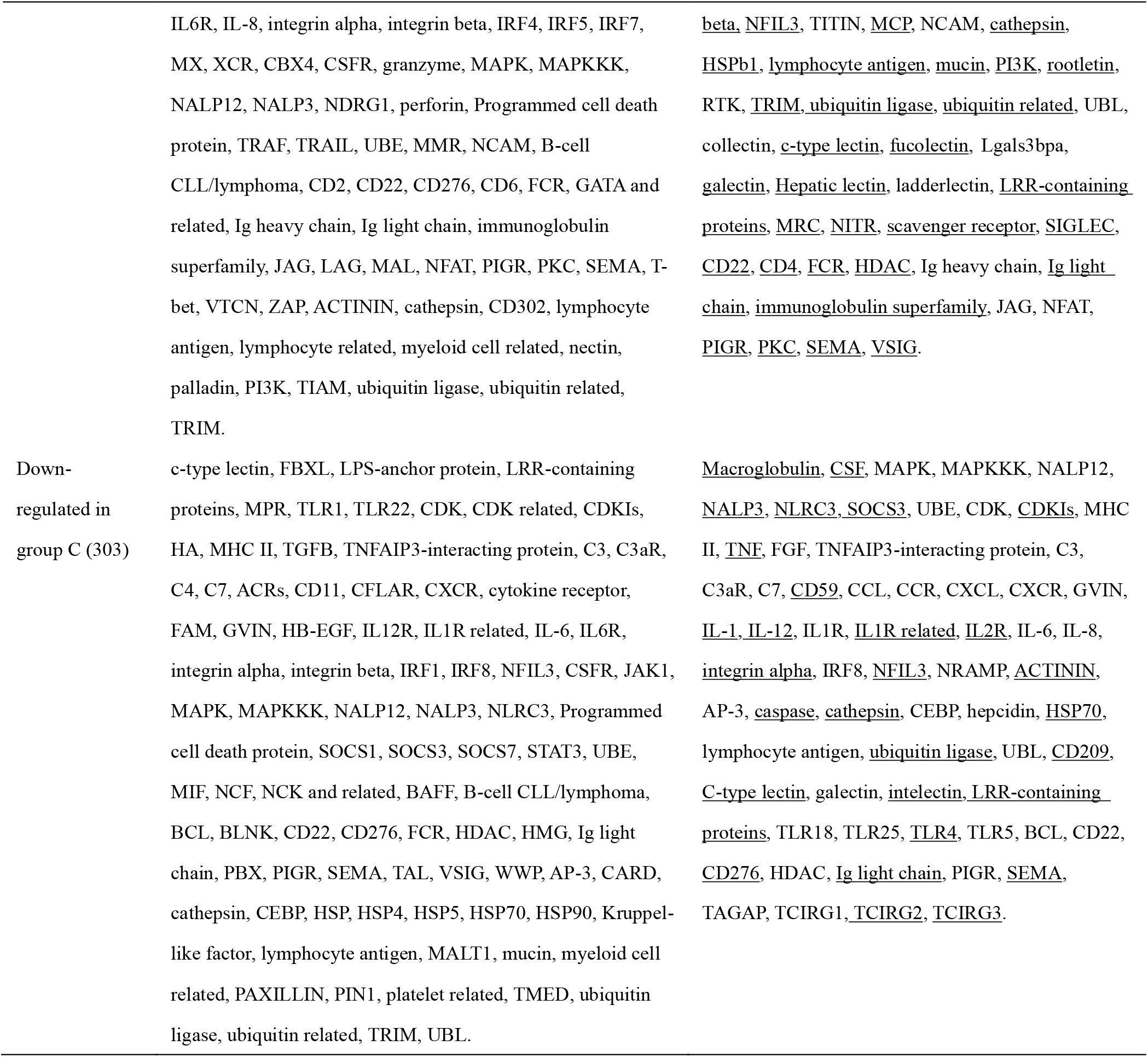
Immune gene categories involved in differentially expressed mRNA for comparison groups A, B and C

**Table 2B.** Immune gene categories involved in differentially expressed mRNA each comparison group D and E

| Compared group | Immune gene category |
| --- | --- |
| Up-regulated in group D (12) | 1<FC<2: <u>MHC I</u>, <u>GVIN</u>, <u>IL21R</u>, <u>NLRC3</u>;<br>2<FC: <u>NALP3</u>, <u>perforin</u>, <u>STAT3</u>, <u>integrin alpha</u>, <u>platelet related</u>, <u>ubiquitin related</u>, <u>MRC</u>, <u>HDAC</u>. |
| Down-regulated in group D (7) | 1<FC<2: <u>TLR18</u>, <u>C3</u>, <u>CCL</u>, <u>integrin beta</u>.<br>2<FC: <u>perforin</u>, <u>IFI</u>, <u>myeloid cell related</u> |
| All in group E (48) | <u>LRR-containing proteins</u>, <u>VEGFR</u>, <u>VEGF</u>, <u>CDKIs</u>, <u>CDK</u>, <u>NFIL3</u>, <u>LIFR</u>, <u>integrin beta</u>, <u>FAM</u>, <u>CCR</u>, <u>CD166</u>, <u>MAPK</u>, <u>CD97</u>, <u>SEMA</u>, <u>PIGR</u>, <u>NFAT</u>, <u>Ig light chain</u>, <u>HMG</u>, <u>HDAC</u>, <u>BTG</u>, <u>BCL</u>, <u>ubiquitin ligase</u>, <u>platelet related</u>, <u>HIF1a</u>, <u>CEBP</u>. |

Notes: the number (>1) of DEG is shown in the brackets. The full names of all gene abbreviations could be found in the Tables S5–S9. The underlined up or down regulated immune gene categories are the regional specific in the Venn diagram for every groups (detailed in Table S10).

### Venn-regional Analysis of Immune-related DEG from Different Comparison Groups

A Venn diagram was created to demonstrate the relationship between the immune-related DEG among groups A-E (Figure 5C). The details for all regions were in Table S10. Though the region-specific DEGs accounted for a large proportion of each comparison group, with immune gene categories for region-specific DEG underlined in Table 2, the overlapping DEGs (Table 3), which contained important genes, were listed with the corresponding immune process and immune gene category in Table 4.

**Table 3.** The immune process and immune gene category for overlapping genes in the Venn regional analysis

| Overlapping regions | Immune process | Immune gene category |
| --- | --- | --- |
| Group A & C (23) | pattern recognition (7), complement system (4), inflammatory cytokines and receptors (3), adapters, effectors and signal transducers (4), innate immune cells related, T/B cell antigen activation (2), other genes related to immune response (2) | CSF, NALP12, UBE, C3, GVIN, IL23R, integrin alpha, NCAM, ubiquitin ligase, C-type lectin, galectin, ladderlectin, LRR-containing proteins, MRC, PIGR |
| Group A & D (1) | inflammatory cytokines and receptors | IFI |
| Group B & C (176) | pattern recognition (18), antigen processing and regulators (14), complement system (4), inflammatory cytokines and receptors (38), adapters, effectors and signal transducers (23), innate immune cells related (8), T/B cell antigen activation (24), other genes related to immune response (47) | Granzyme, MAPK, MAPKKK, NALP12, NALP3, NLRC3, perforin, Programmed cell death protein, SOCS1, SOCS3, SOCS7, UBE, CDK, CDK related, CDKIs, HA, MHC II, TGFB, TNF, TNFAIP3-interacting protein, VEGF, C3aR, C7, ACRs, CCL, CCR, CFLAR, CXCL, CXCR, cytokine receptor, FGFR, GVIN, IL1, IL17R, IL1R related, IL2R, IL6, IL6R, IL8, integrin alpha, IRF8, NFIL3, TITIN, MIF, NCAM, NCF, NCK and related, NRAMP, ACTININ, AP-3, CARD, cathepsin, CEBP, hepcidin, HSP, HSP5, HSP70, lymphocyte antigen, paladin, PIN1, platelet related, RTK, TMED, TRIM, ubiquitin ligase, ubiquitin related, UBL, collectin, C-type lectin, FBXL, galectin, LRR-containing proteins, MPR, RBL, TLR1, TLR13, TLR22, TLR25, TLR5, BCL, BLNK, CD22, FCR, HDAC, HMG, Ig heavy chain, Ig light chain, JAG, NFAT, PBX, SEMA, TAGAP, TCIRG1 |
| Group B & E (4) | antigen processing and regulators (3), other genes related to immune response | CEBP, VEGFR |
| Group B & D (1) | inflammatory cytokines and receptors | integrin alpha |
| Group C & D (1) | inflammatory cytokines and receptors | CCL |
| Group A & B & C (5) | pattern recognition, inflammatory cytokines and receptors (3), adapters, effectors and signal transducers | LRR-containing proteins, CXCL, IL1R, perforin |
| Group A & B & D (1) | complement system | C3 |
| Group B & C & D (2) | adapters, effectors and signal transducers (2) | perforin |
| Group A & B & C & D (2) | pattern recognition, inflammatory cytokines and receptors, | TLR18, IL21R |

### Immune related GO Terms as well as Network Correlation between LncRNAs and Genes in Immune Related Module

The results of “WGCNA” were selected to determine the biological processes involved in CyHV-3-induced modulation of genes by lncRNAs, from an immune perspective. Among the currently divided 35 gene modules (Figure 6A), 12 modules were found with immune-related GO terms (Figure S1). Generally, “double-stranded DNA binding”, “receptor-mediated endocytosis”, “response to bacterium”, “lysosome”, “defense response to fungus”, “killing of cells of other organism”, “oxygen carrier activity”, “oxygen binding”, “defense response to bacterium”, “killing of cells of other organism”, “NuA4 histone acetyltransferase complex”, “defense response to gram-negative bacterium”, “defense response to gram-positive bacterium”, “immune response”, “regulation of cytokine secretion”, “cytolysis”, “execution phase of apoptosis”, “cellular response to gamma radiation”, “nucleic acid metabolic process”, “TOR signaling”, “DNA replication”, “ubiquitin-dependent protein catabolic process”, “granulocyte differentiation”, and “macrophage differentiation” were revealed after the “WGCNA” analysis. The top five immune-related modules containing most immune-related GO terms (Figure 6B) were analyzed for the relationship between LncRNA and transcripts by PPI networks in cystoscope (Figure 6C, with details in Table S11). The LncRNA regulated genes, which upregulated in group C, such as Semaphorin-4E in the blue module, can be the factor associated with the resistance in the breeding strain. The LncRNA regulated genes, which downregulated in group C, such as Lgals3l in the grey60 module, can be the factor associated with less accessible to CyHV-3 in the breeding strain. The LncRNA regulated genes, which was upregulated in group B and downregulated in group C, such as natural resistance-associated macrophage protein (Nramp), plasminogen activator inhibitor (PAI) in the brown module, could be the key clues for susceptibility of CyHV-3 in the non-breeding strain. Specifically, for detailed correlations of lncRNAs and DEG in groups A to C (Table S12), T cell leukemia homeobox 3 (TLX3) and LGALS3 were revealed, respectively, in the survivors of the breeding strain.

**Figure 6.**
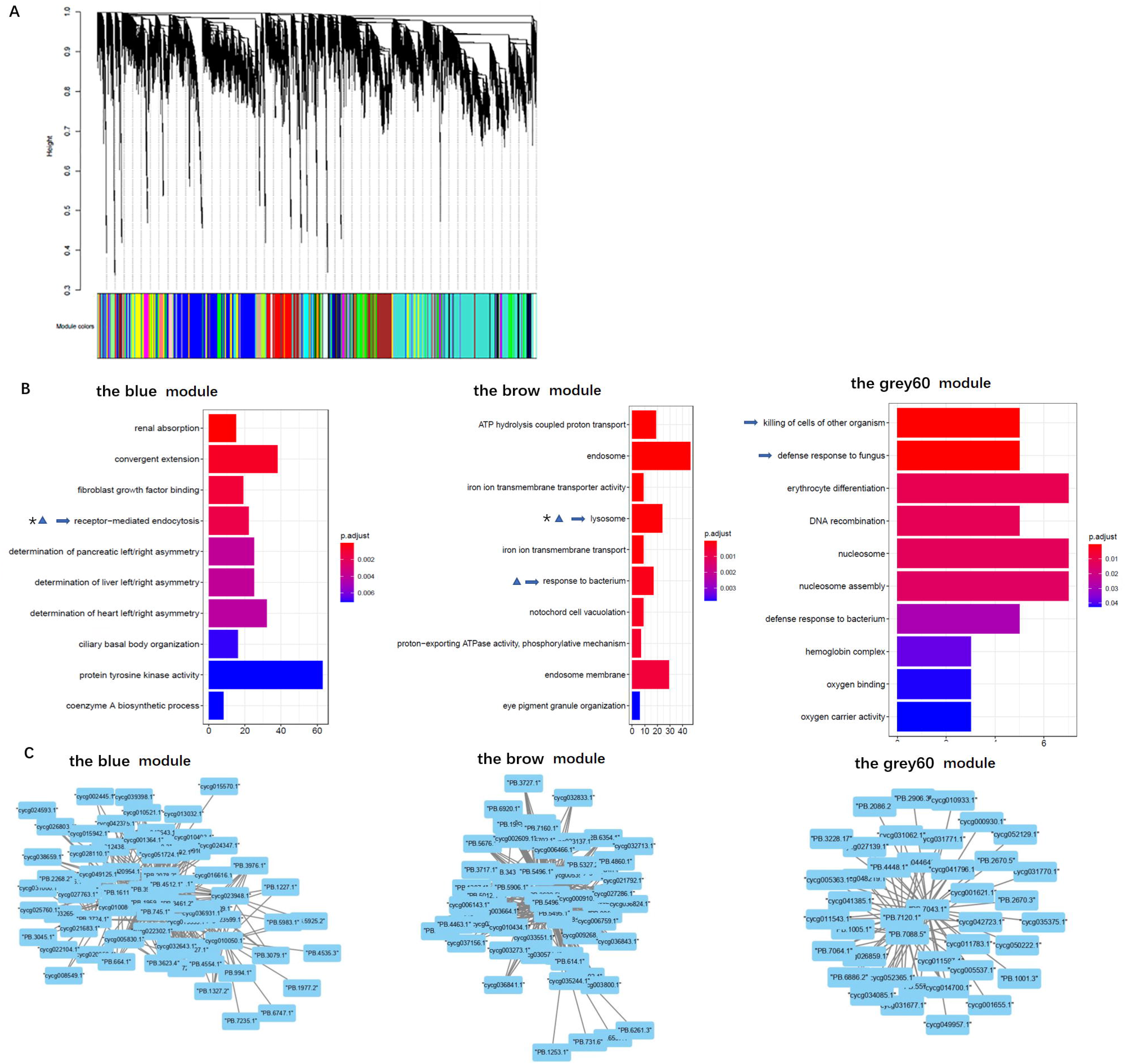
Gene ontology analysis and cystoscopes of key matched lncRNA and transcripts in the immune related module. (A) the cluster dendrogram of all gene module. (B) The GO terms involved in immune-related module, including the blue, brow, and the grey60 module, which contain the matched lncRNA and transcripts involved in current analyzed comparison groups. The star, triangle, and arrow represent the revealed GO term for comparison group A, B, and C, respectively. **(C)** cystoscopes of key matched lncRNA and transcripts in the immune related module, as shown for blue, brow and grey60 module, respectively.

## DISCUSSION

This study demonstrated the anti-CyHV-3 immune mechanisms of a breeding strain of common carp. Current revealed dramatically decreased mortality and less inner tissues swelling together with the previously revealed reduced virus load of tissues (16, 17), proved the resistance to CyHV-3. Current found biological processes and pathways by transcriptomic analysis has been merged to improve clarity. Afterwards, by comparing the survivors and healthy fish in either breeding or non-breeding strains, key genes and related lncRNAs involved in immune processes were also revealed. Accordingly, the immunogenetically insensitive or susceptible factors to CyHV-3 infection were determined.

Based on data of the healthy state, the current results suggested potential components involved in the entry of the virus. Firstly, integrin acts as a herpesvirus receptor (29), and the integrin-dependent signalosome in herpesvirus-infected cells mediated or coactivated numerous inflammatory responses and signaling transductions (29). Therefore, the current finding that integrin beta 1 (ITGB1) was significantly downregulated in the breeding strain compared with the non-breeding strain, even during the steady status, may indicate that ITGB1 facilitates the binding of herpes virus glycoprotein for entry (30). This coincides with the finding that integrin signaling promotes the release of intracellular calcium stores and contributes to viral entry and cell-to-cell spreading via glycoprotein H, during herpes simplex virus infection in humans (31). Recent reports have also demonstrated that some integrins on lymphocytes, such as B cells, could facilitate the mucosa-specific homing (32). Secondly, higher TLR expression in survivors of the non-breeding strain compared with that of the breeding strain also indicated its role in virus entry and downstream proinflammatory signaling. Of note, TLR4 signaling leads to the production of proinflammatory cytokines in human lymphatic endothelial cells (29). In addition, fish-specific and virus-responding TLR18 (33, 34) (the most overlapping gene among groups) was downregulated in survivors of the breeding strain, whereas it was upregulated in those of the non-breeding stain. This suggested its potential ability to facilitate CyHV-3 infection. MAPK signaling, as the downstream process of pattern recognition receptors, could facilitate the tumor necrosis factor-alpha (TNFA)-induced suppressor of cytokine signaling 3 (SOCS3) expression. This can lead to both pro-inflammatory immune response and failure in growth (35), according to the upregulated genes upon infection observed in the non-breeding strain in the comparison group B.

Further, by comparing the survivors from different stains, DEGs of the inflammatory status also provide clues for how to block virus. Non-specific binding of the virus by lectins played a protective role in preventing virus entry. Both upregulated ladderlectins and higher galectin expression were detected in survivors of breeding strains compared with non-breeding strains. This is suggestive of their blocking ability for viral proteins (36, 37), such as glycoprotein (38). Moreover, as the head kidney is one of the major reticuloendothelial systems in fish (39), mucin was also found to be upregulated as the anti-virus barrier. This was shown in the comparison group B as the gel-forming mucin 5B (MUC5B) (40) was upregulated in the survivors of the non-breeding strain. However, higher expression levels of membrane-bound MUC3 (41) were found in survivors of the breeding strain versus the non-breeding strain. Additionally, in the comparison group C, which comparing the survivors from both strains, the upregulated chemokine CCL4 homologues, may indicate that CCL4 in the breeding strain may bind the virus and be a restrictor of virus infection, as the reported function in human (42). These findings suggested that more mucus was secreted, thereby causing tissue swelling upon CyHV-3 infection in the non-breeding strain. For the survivors of the breeding strain, the membrane-bound mucin could effectively bind the virus with no gel.

Lectins are involved in the clearance of apoptotic cells via the complement pathway. Hence, in survivors, the complement components reported in the acute phase in common carp during CyHV-3 infection (43), could facilitate the phagocytic process via binding of MRC (Mannose-Receptor C) in fish (44). In the comparison of the survivors from both strains for group C, the observed difference in the phagocytic process suggested the clearance of apoptosis cells, which was indicated by the upregulated ubiquitin ligase in the survivors of breeding strain. Meanwhile, IL-1, IL-6, and IL-8 were upregulated in group B. With regard to cytokine signaling, there were less proinflammatory cytokines and immune regulation-related cytokine receptors upregulated in the survivors of the breeding strain compared with the non-breeding strain. The interleukin 21 receptor (IL21R), which was found regulated in the survivors of the breeding strain for group A and the healthy control of the non-breeding strain for group B, indicated the control of inflammation by suppressing STAT3 signaling (45). The upregulated IL23R in the survivors of the breeding strain for group A can promote cytotoxic ability (46), which may immediately kill infected cells. The lack of antigen presentation in comparison group A suggested that the infection was overcome before the amplification of CyHV-3 in the breeding strain. In contrast, in group B, numerous upregulated genes were involved in antigen presentation in the non-breeding strain. Therefore, the survivors from breeding strains overcame the infection mainly through phagocytosis and cytotoxicity at the cellular level, without activating the typical process of proinflammation as in survivors from the non-breeding strain.

Thus, there are different signatures between the two strains, regarding to the survival strategy. For the survivors in the breeding strain, self-repairing related autophagy was detected as both the KEGG pathways “regulation of autophagy” and autophagy-related fish antiviral tripartite motif (TRIM) protein (47) were found in group A. These findings were in line with TRIMs, which were found regulated in the survivors of breeding strain for group A and in the healthy control of non-breeding strain for group B. The above finding of TRIM is coincident with one of recent revealed CyHV-3 resistant related DEGs with identified QTLs (13). In the survivors from the breeding strain, the higher PI3K also suggests higher autophagy, since the PI3K/AKT/mTOR pathway enhances this process (48). Also, in the breeding strain the suppressed IFN activation was also suggested, for that fish TRIM may inhibit the activation of IFN and attenuate IFN regulatory factor (IRF) (49, 50). The factor that there was more expression of TRAF6 in survivors of breeding strain compared with those of non-breeding strain, is in line with the resistant related SNP on TRAF6 (51), and suggests the possible repression on production of type I IFN (52). In addition, nuclear factor, interleukin 3 regulated (NFIL3) can control Treg cell function via directly binding to and negatively regulates the expression of Foxp3 (53), and has been revealed stimulating both proinflammatory (e.g., NF-kappa B [NF-κB]) and anti-inflammatory factors (e.g., IL10) in carp (54). Thus, the upregulated NFIL3 in the survivors of the breeding strain for group A may suggested the extensive activation of immune cells, with diminished immune regulation. The directed lymphocytes response could be present as there was upregulation of IL23R, which significantly enhances the expression of cytotoxic mediators (46), as well as cathepsin L (a component of lysosomes) (55). This indicated an enhanced activation of the cytotoxic ability of T cells in survivors from the breeding strain. For B cells, the secretion of Ig-related genes (e.g., polymeric immunoglobulin receptor [PIGR], Ig heavy chain and light chain) was also upregulated in the survivors of breeding strain for group A. To fight the virus, the survivors from the non-breeding strain developed typical inflammatory cascades, including pro-inflammatory cytokines (e.g., IL-1, IL-6, and IL-8), as well as MAPK signaling, which are profoundly involved in cell survival functions during viral infection (56). For the control of inflammation, the survivors from the common strain exhibited suppression of the IFN response, which was also reflected by the upregulated IRF8, the inhibitor of the MYD88-mediated NF-κB signaling pathway (57). Downstream hypoxia was also found, as indicated by the “p53 signaling pathway” and “oxidative phosphorylation”. The hypoxic status could also protect survivors of the non-breeding strain from death, since p53 suppresses cell apoptosis (58), and HIF1A regulates virus-induced glycolytic genes (59).

Furthermore, the advantages of disease-resistant genes were reflected by comparing the survivors from the two strains in comparison group C, which can provide clues for how to develop resistance to CyHV-3. There was more semaphorin, which is related to immunoregulation (60, 61), in survivors from breeding strains than non-breeding strains, and was also regulated by LncRNA. This indicated a possible tolerance of viral replication or latency after a primary infection in the survivors (2), while had no negative effects on the proliferation of host cells (62). There was less cyclin-dependent kinase inhibitor 1D (CDKN1D) in survivors from the breeding strain compared with the non-breeding strain. This indicated that there was an interrupted circadian cell-cycle timing in the survivors from non-breeding strains (63). The higher levels of STAT3 and possible its upregulated integrins (64) in survivors from the non-breeding strain may facilitate the effect of enhanced IL6 signaling. Apart from the typical immune signals in comparison group C, there was relatively higher PI3K activity, which was involved in the KEGG pathway “dopaminergic synapse”. This indicated that dopamine inhibited inflammation (65) in survivors from the breeding strain. This is because PI3K is dependent on the accumulation of DOPA decarboxylase, the enzyme involved in the production of dopamine, which is reduced by a viral infection (66). Additionally, genes responsive to the secretion of immunoglobulin (e.g., PIGR and Ig light chain), which is important for anti-virus immunity, were common (in comparison group E) between the two strains.

The comparison of two strains with healthy status in group D suggested a basal anti-virus immunity. Among the upregulated genes in the healthy control of breeding strain for comparison group D, the expression of NACHT, LRR, and PYD domains-containing protein 3 (NALP3) indicated a stronger ability to form the inflammasome in the breeding strain. The higher levels of NLR family CARD domain containing 3 (NLRC3), which is a negative regulator of the DNA sensor STING, suggest less sensitivity to the DNA virus in the breeding strain. The higher levels of MHC I indicate greater potential to activate response by cytotoxic CD8+ T cells upon herpes virus infection, as previously revealed in the resistant strain R3 (67). For the potential T helper cell differentiation-related JAK/STAT pathway, the upregulation of both STAT3 and histone deacetylase (HDAC) indicated an easier differentiation of both T helper 17 and regulatory T directions (68), respectively, in the breeding strain even in the steady state (69, 70). Upregulation of HDAC in the healthy control of breeding strain for group D, that could inhibit the function of macrophages in fish (71), also suggests a more delicate immune regulation during the steady state in the breeding strain. However, among the downregulated genes, which indicated higher expression in the non-breeding stain at a steady state, TLR18 and ITGB1 suggested susceptibility to virus binding. Moreover, C-C motif chemokine ligand 4 (CCL4), which can protect infected cells with viral burden (72), may be a risk factor for the non-breeding strain.

The revealed regulation of resistance to CyHV-3 by lncRNA involved numerous biological processes. Among the revealed GO terms in comparison group B, “DNA replication” could possibly indicate virus proliferation in the non-breeding strain. In survivors from the non-breeding strain, the lncRNA-regulated genes were mainly involved in innate immune cell function (e.g., macrophage protein and cathepsin, as components of lysosomes) and cell apoptosis. Nramp and PAI were found regulated by LncRNA in comparison groups B (up) and C (down). This finding indicates the more expression of Nramp and PAI in the survivors of non-breeding strain, could facilitate virus infection and proliferation for infected cell respectively, since that Nramp may serve as virus receptor (73), meanwhile PAI can inhibits apoptosis in cell lines infected with viruses (74). While, in survivors from the breeding strain, the only lncRNA-modulated gene, TLX3, has been shown to be strongly methylated. This indicates that TLX3 expression was suppressed during hepatitis B virus-related cancer (75). This suggests that TLX3 in survivors from the breeding strain may play a protective role, and participate in lymphocyte proliferation in the head kidney. Meanwhile, echoing with recent finding of the participation of zebrafish galectin proteins in immunity against viral infection (38, 76), in comparison group C the LncRNA regulated transcripts Lgals3l (down) and Lgals3bpa (up), as well as the upregulated galectin 3 suggested the regulation of galectin-3 related biological activities could be related to reduce the viral attachment for survivors of the breeding strain.

Therefore, based on the present findings, a hypothesis has been proposed for the immune mechanisms involved in both healthy controls and survivors from infection in both strains (Figure 7). In conclusion, the breeding strain of common carp showed a better ability to maintain immune homeostasis in both steady and inflammatory states, and displayed enhanced blockage of CyHV-3 infection compared with the non-breeding strain. Thus, this strain could be termed as a resistant strain accordingly. Since the modulation of mRNA and lncRNA expression dynamics currently remains a hypothesis, further molecular evidence is needed to elucidate the mechanism of both resistance and susceptible. In addition, the finding that both the inhibition of inflammation by dopamine in the breeding strain and the disrupted bio-clock in the non-breeding strain upon CyHV-3 infection suggested better growth performance for the breeding strain. Therefore, the possible advantage in this resistant strain for growth performance warrants further study.

**Figure 7.**
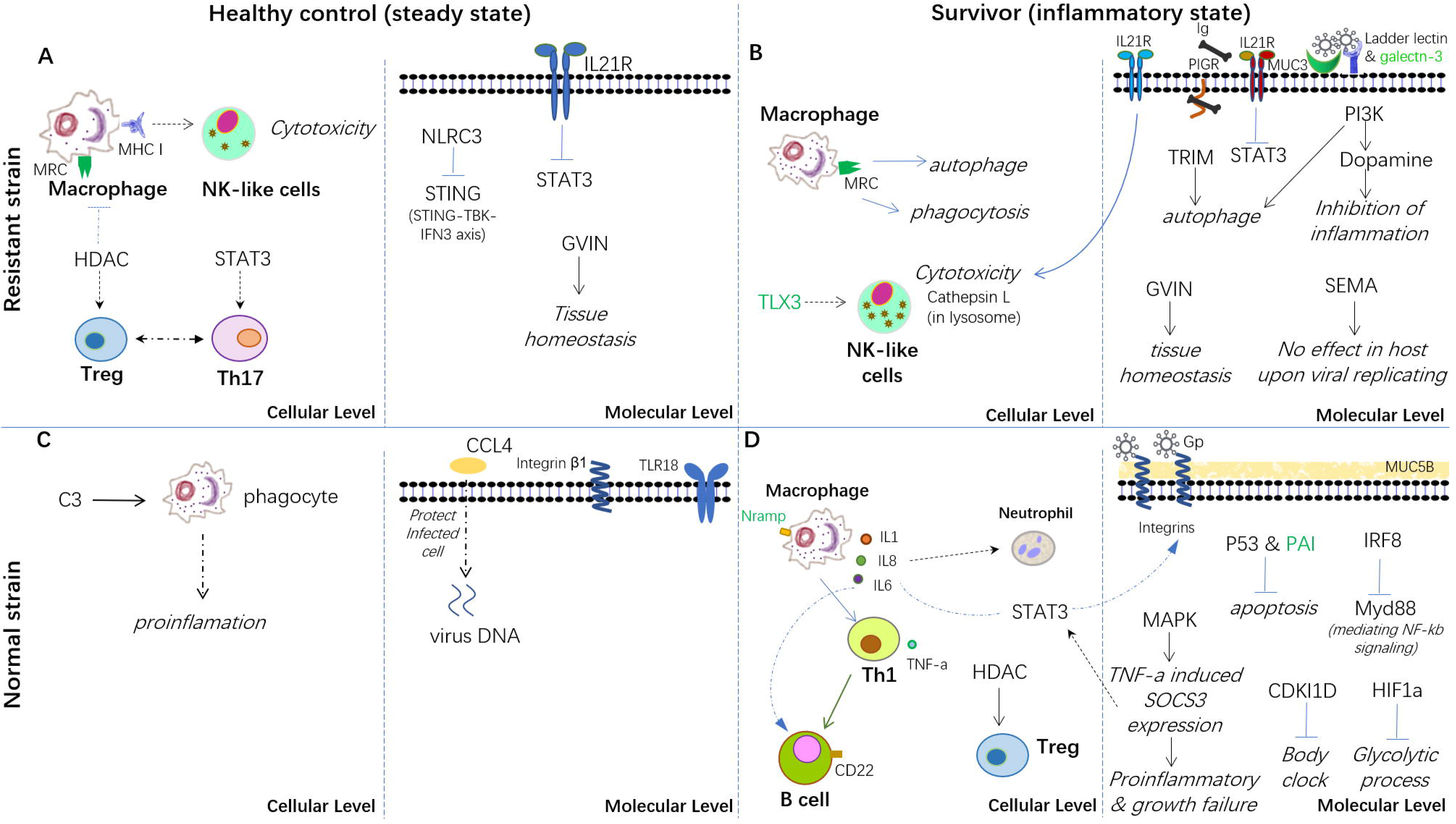
Hypothesis of the carp immune mechanism in both breeding (resistant) and non-breeding (normal) strains, either at the steady state or upon surviving for CyHV-3 infection. (A) healthy control in the steady state in the breeding strain. (B) survivors in an inflammatory state in the breeding strain. (C) healthy control in the steady state in non-breeding strain. (D) survivors in an inflammatory state in the non-breeding strain. The words in italics, bold, and green represent the biological process, cell type, and lncRNA-modulated immune genes, respectively. The arrow denotes facilitation or promotion, and ---| denotes inhibition. The dashed line indicates a possible correlation.

## AUTHOR CONTRIBUTIONS

Lianyu Shi and Zhiying Jia conceived the project and designed the experiments. Nan Wu and Zhying Jia wrote the manuscript. Zhying Jia performed the experiments. Zhying Jia and Jiaxin Sun conducted the CyHV-3 infection experiment and collected samples. Chitao Li, Xuesong Hu and Yanlong Ge performed fish propagation and culture. Xiaona Jiang conducted the RT-PCR experiment. Nan Wu and Xiao-Qin Xia coordinated the data analysis tasks. Nan Wu, Heng Li, Mijuan Shi, Xiaona Jiang, Weidong Ye, Ying Tang, and Yingyin Cheng analyzed the data. The manuscript was revised and approved by Xiao-Qin Xia and Lianyu Shi.

## ACKNOWLEDGMENTS

This work was funded by grants from the National Key R&D Program of China (2018YFD0900302-6), The Natural Science Foundation of Heilongjiang Province (TD2019C004), China Aquaculture Research System (CARS-45-06), and Central Public-interest Scientific Institutional Basal Research Fund (2020TD31). We also thank Mr. Bruno Unger from University of Canterbury in New Zealand for language editing.

## CONTRIBUTION TO THE FIELD STATEMENT

Cyprinid herpesvirus-3 (CyHV-3) is the major threat to common carp aquaculture. Experimental infections of carp from pure lines or crosses have indicated the existence of genetic background of resistance by divergent survival rates. The immune response of carp to CyHV-3 involves both innate and adaptive aspects. Even non-coding RNAs play an important role in anti-CyHV-3 immunity. However, there is a lack of systemic studies focusing on a detailed network of the immune system for the anti-CyHV-3 immune mechanism in common carp. Benefiting from a strain of common carp which has already shown a higher survival rate in CyHV-3 infection, this study systematically illustrated the immune mechanisms involved in anti-CyHV-3 ability on both mRNA and non-coding RNA levels, demonstrated that the carp may obtain significant resistance to CyHV-3 through a specific innate immune mechanism, and revealed the involved biological processes including autophagy, phagocytosis, cytotoxicity, blocking of the virus by lectins and MUC 3, and the immune suppressive signal transduction. The immune factors involved in resistant or susceptible to the disease shed light on prevention of CyHV-3 infection genetically.

## DATA AVAILABILITY STATEMENT

The datasets, containing all clean data of this transcriptomic analysis, can be found in the Genome Sequence Archive (GSA) database (http://gsa.big.ac.cn/index.jsp) with the BioProject identifier <CRA003105>.

