## Supplemental figure for "Integrative Transcriptomic Analysis Reveals the Immune Mechanism for A CyHV-3-Resistant Common Carp Strain"

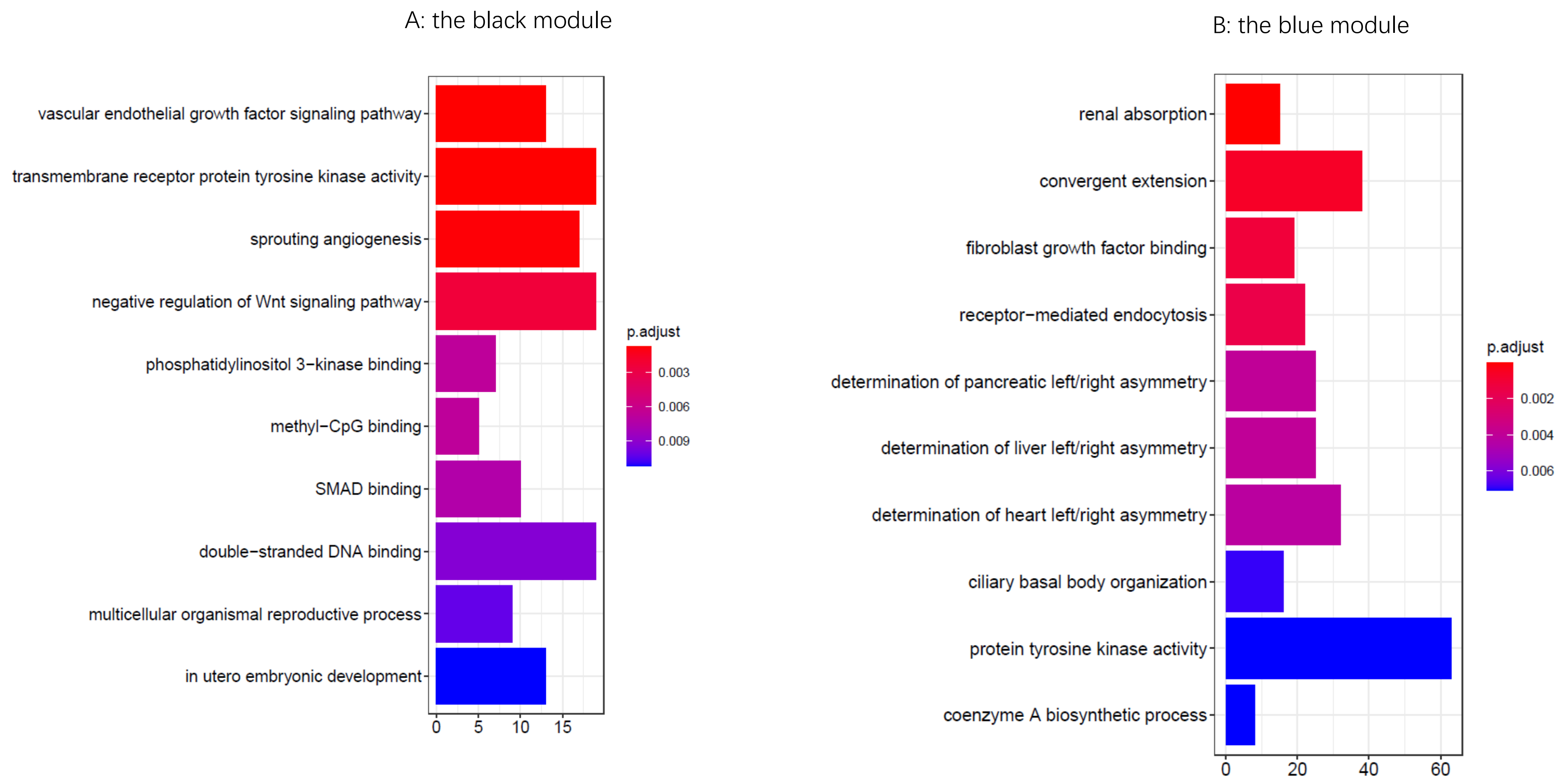
A: the black module
vascular endothelial growth factor signaling pathway
transmembrane receptor protein tyrosine kinase activity
sprouting angiogenesis
negative regulation of Wnt signaling pathway
phosphatidylinositol 3−kinase binding
methyl−CpG binding
SMAD binding
double−stranded DNA binding
multicellular organismal reproductive process
in utero embryonic development
0
5
10
15
p.adjust
0.003
0.006
0.009
B: the blue module
renal absorption
convergent extension
fibroblast growth factor binding
receptor−mediated endocytosis
determination of pancreatic left/right asymmetry
determination of liver left/right asymmetry
determination of heart left/right asymmetry
ciliary basal body organization
protein tyrosine kinase activity
coenzyme A biosynthetic process
0
20
40
60
p.adjust
0.002
0.004
0.006

C: the brown module

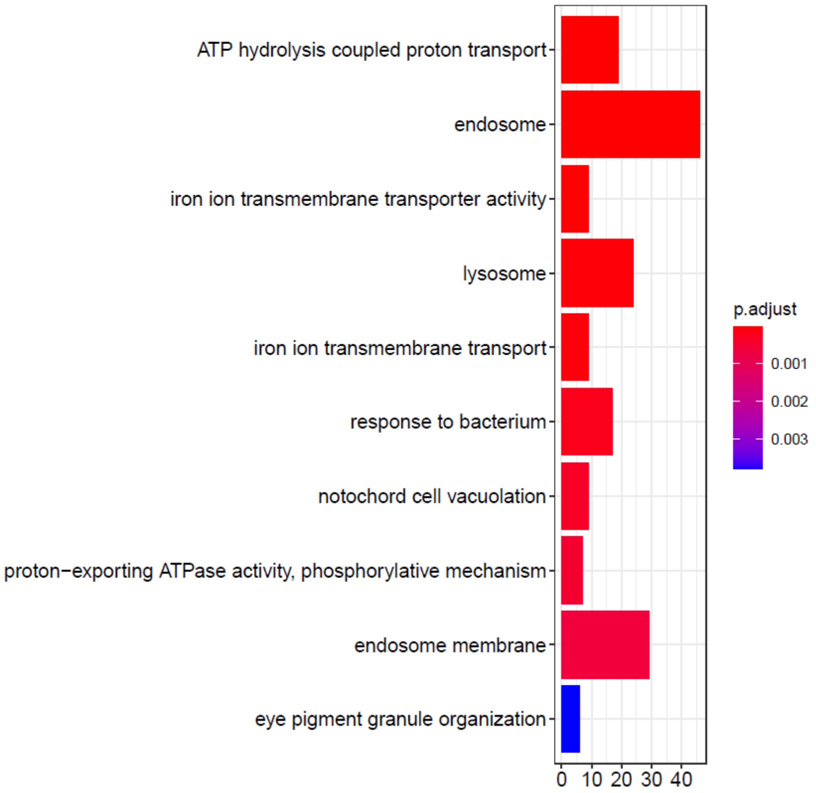


D: the dark orange module

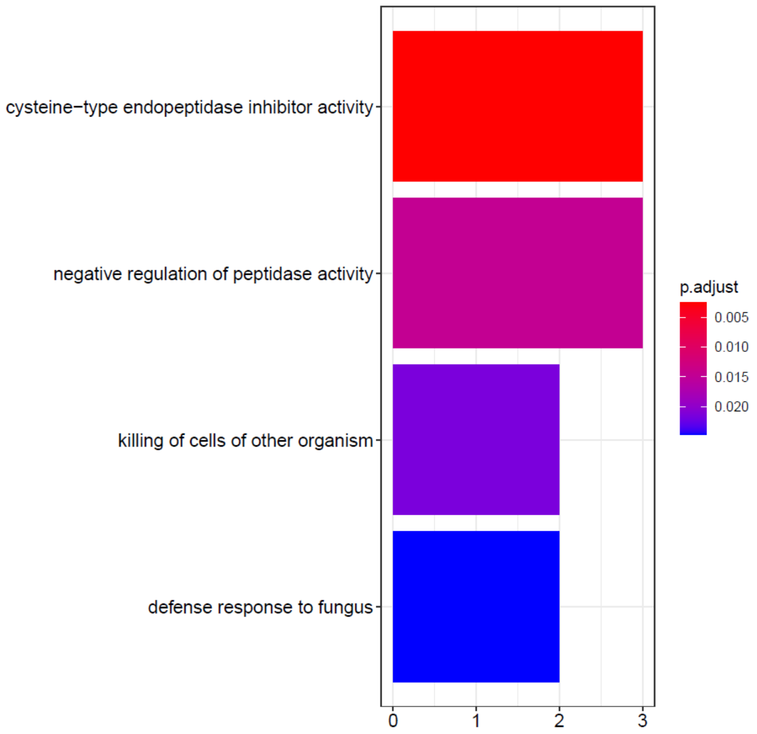

E: the grey 60 module

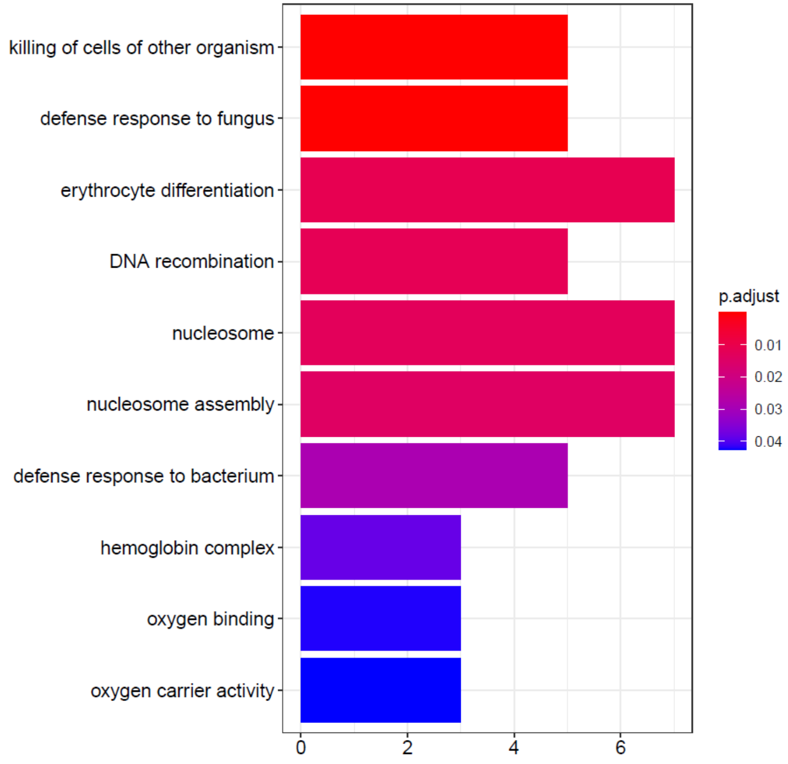


F: the light green module

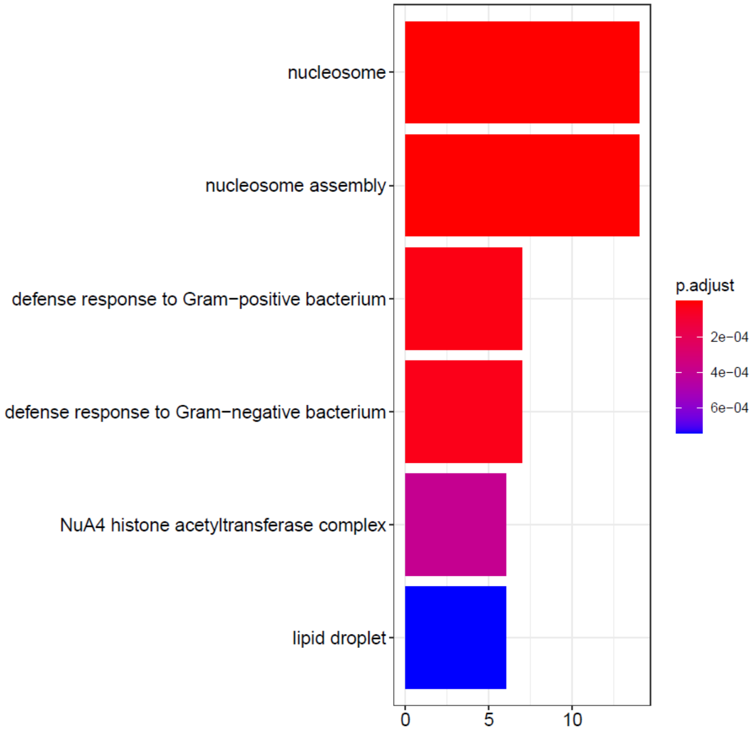

G: the light yellow module

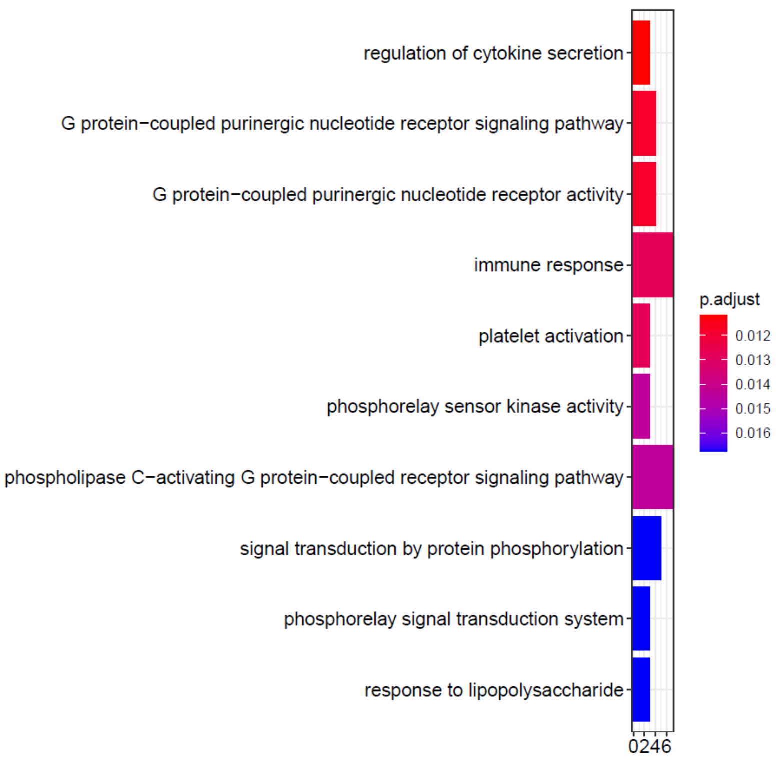


H: the midnight blue module

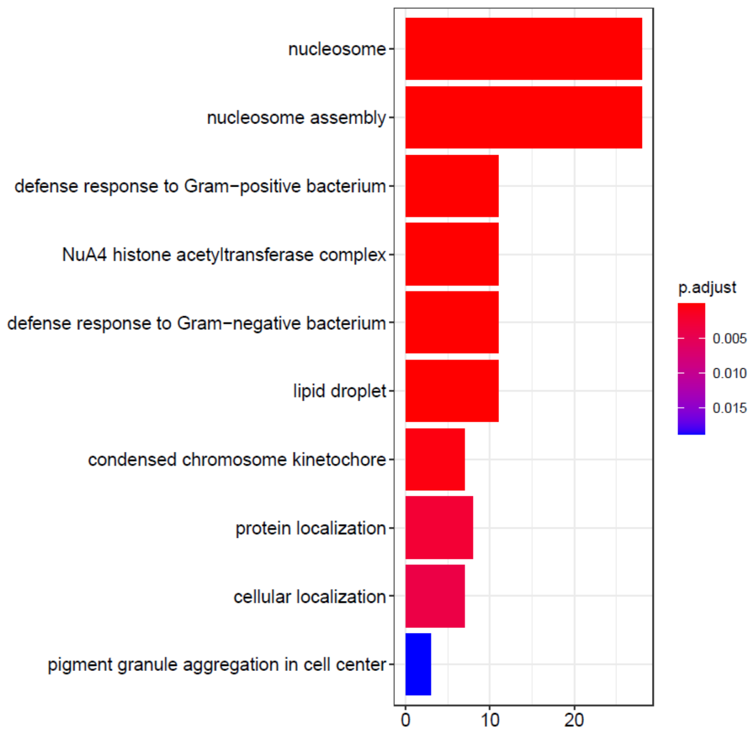

I: the orange module

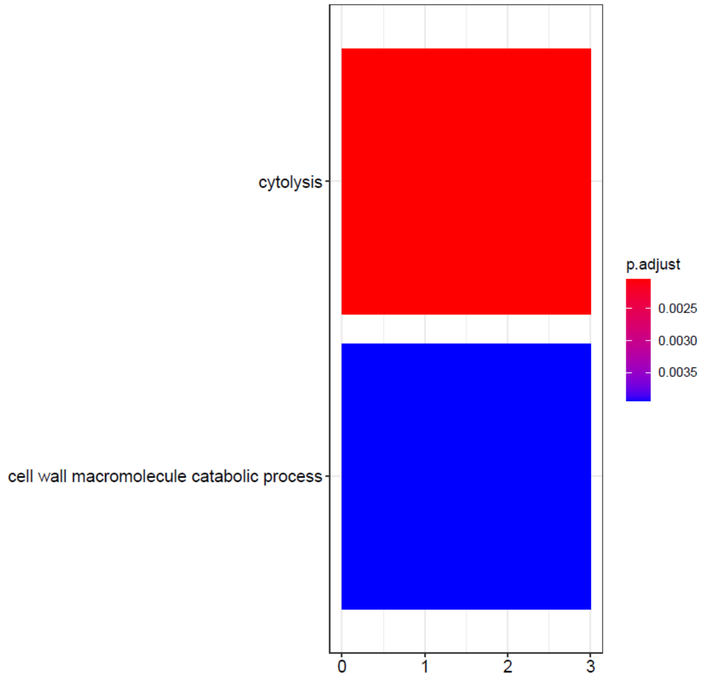


J: the pink module

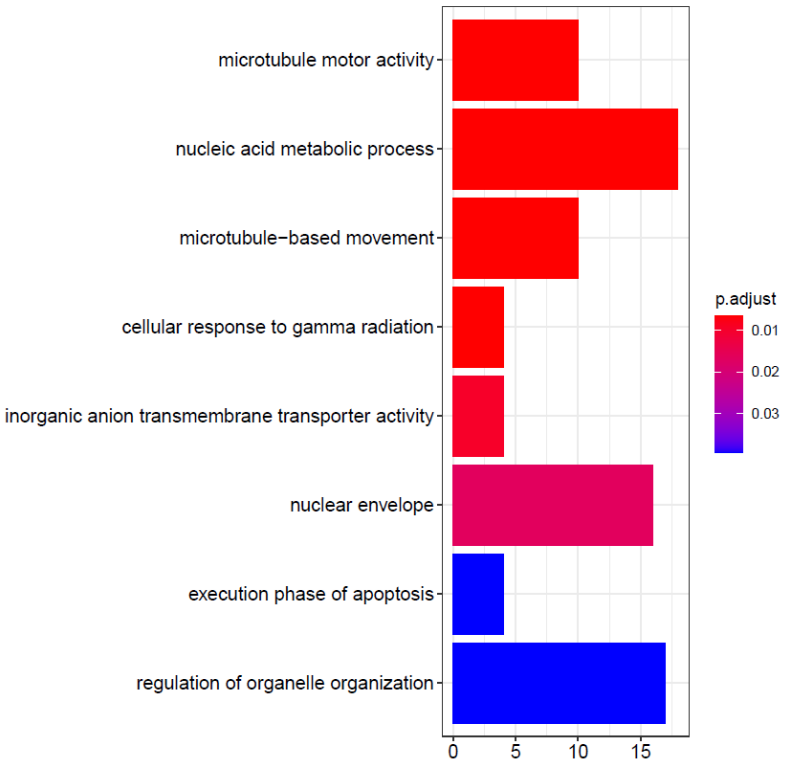

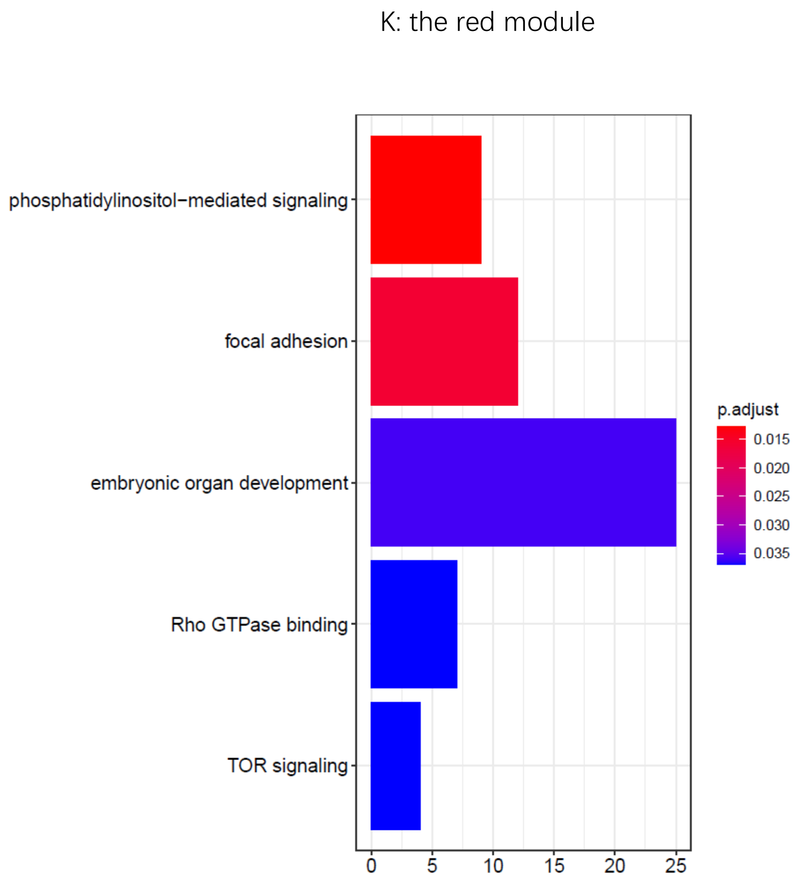
K: the red module
phosphatidylinositol−mediated signaling
focal adhesion
embryonic organ development
Rho GTPase binding
TOR signaling
0
5
10
15
20
25
p.adjust
0.015
0.020
0.025
0.030
0.035

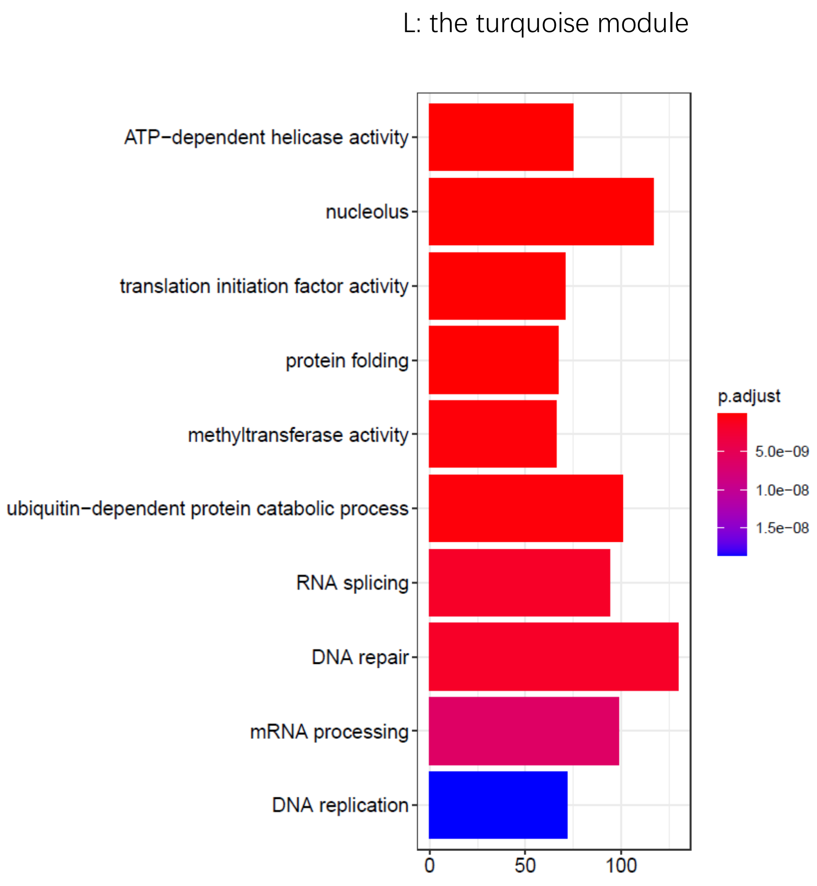
L: the turquoise module
ATP−dependent helicase activity
nucleolus
translation initiation factor activity
protein folding
methyltransferase activity
ubiquitin−dependent protein catabolic process
RNA splicing
DNA repair
mRNA processing
DNA replication
0
50
100
p.adjust
5.0e−09
1.0e−08
1.5e−08

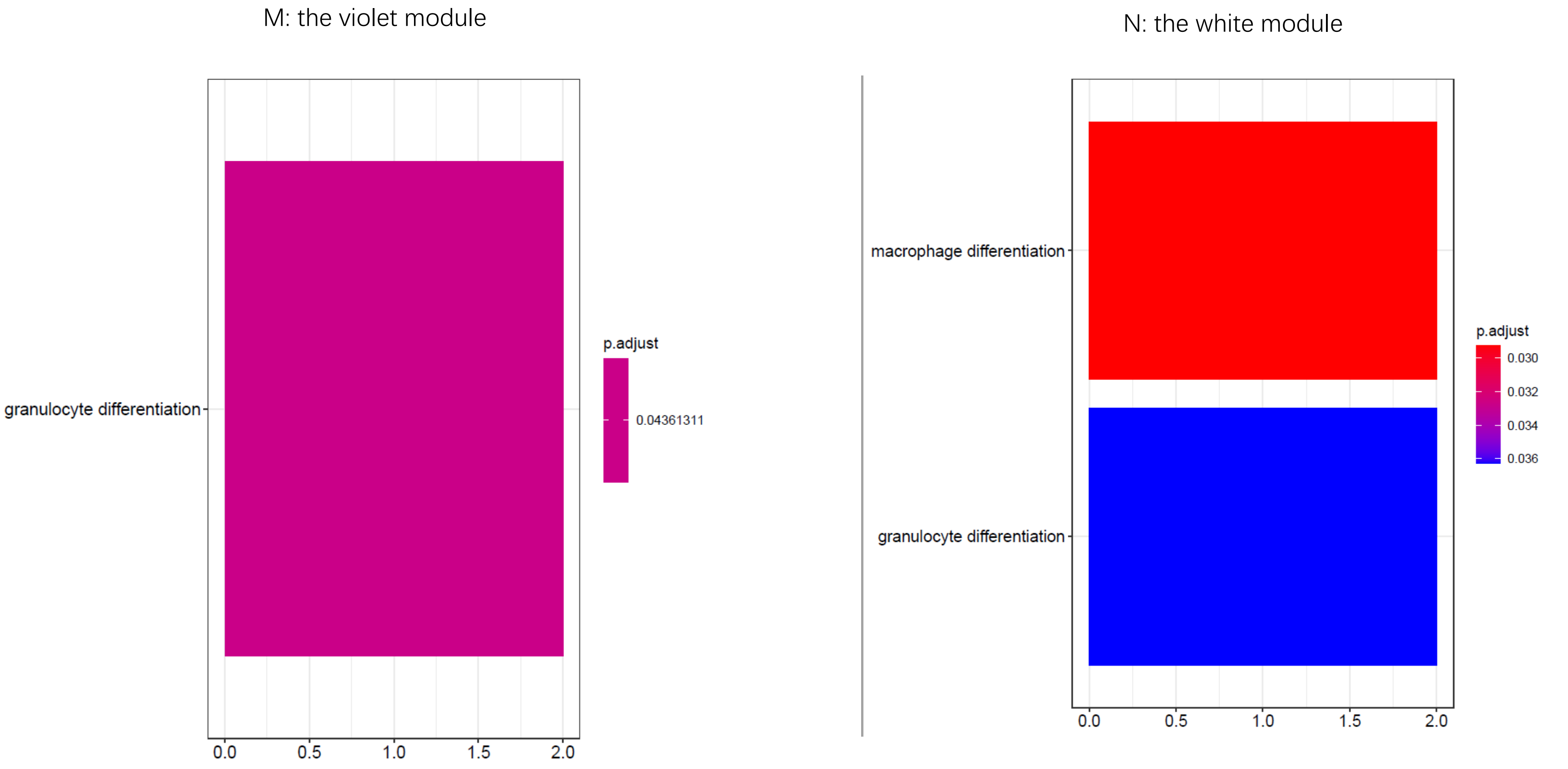


Fig. S1 the immune go terms containing module
